# Controlled Release of Anti-inflammatory and Pro-angiogenic Factors from Macroporous Scaffolds

**DOI:** 10.1101/2020.09.25.314344

**Authors:** Jia-Pu Liang, Robert P. Accolla, Kaiyuan Jiang, Ying Li, Cherie L. Stabler

## Abstract

The simultaneous local delivery of anti-inflammatory and pro-angiogenic agents via biomaterial scaffolds presents a promising method for improving the engraftment of tissue-engineered implants while avoiding potentially detrimental systemic delivery. In this study, PDMS microbeads were loaded with either anti-inflammatory dexamethasone (Dex) or pro-angiogenic 17β-estradiol (E2) and subsequently integrated into a single macroporous scaffold to create a controlled, dual drug-delivery platform. Compared to a standard monolithic drug dispersion scaffold, macroporous scaffolds containing drug-loaded microbeads exhibited reduced initial burst release and increased the durability of drug release for both agents. Incubation of scaffolds with LPS-stimulated M1 macrophages found that Dex suppressed the production of pro-inflammatory and pro-angiogenic factors, when compared to drug-free control scaffolds; however, the co-incubation of macrophages with Dex and E2 scaffolds restored their pro-angiogenic features. Following implantation, Dex-loaded microbead scaffolds (Dex-µBS) suppressed host cell infiltration and integration, when compared to controls. In contrast, the co-delivery of dexamethasone with estrogen from the microbead scaffold (Dex/E2-µBS) dampened overall host cell infiltration but restored graft vascularization. These results demonstrate the utility of a microbead scaffold approach for the controlled, tailored, and local release of multiple drugs from an open framework implant. It further highlights the complementary impacts of local Dex and E2 delivery to direct the healthy integration of implants, which has broad applications to the field of tissue engineering and regenerative medicine.

## 1. Introduction

Biomaterials are broadly used in regenerative medicine to support cellular transplants and heal defective tissues. Host foreign body responses (FBR) to these grafts, however, present a significant obstacle to their utility. The incorporation of porosity within biomaterial implants can mitigate fibrotic encapsulation, with parameters such as pore size and interconnectivity directing host responses towards integrative features such as vascularization and healthy extracellular matrix deposition (1-3). Immune stimuli associated with surgical injury and the presence of foreign material, however, are largely unavoidable, resulting in transient inflammatory cytokines production that instigates cellular necrosis and negatively impacts graft outcomes (4, 5). The integration of anti-inflammatory agents into these optimized biomaterial scaffolds could serve to locally alleviate these deleterious responses.

Anti-inflammatory glucocorticoids, such as dexamethasone (Dex), are commonly incorporated into biomaterial platforms due to their pleiotropic and potent effects on the implant microenvironment (6-8). Dex inhibits pro-inflammatory cytokine expression in leukocytes, including macrophages, while promoting regulatory macrophage phenotypes (9-11); however, Dex can also globally suppress cellular infiltration and vascular endothelial growth factor (VEGF) expression on host cells, resulting in delayed angiogenesis (11, 12). We observed this dose-dependent modulation on host cell mobility and phenotype following the implantation of Dex-releasing polydimethylsiloxane (PDMS)-based monolithic macroporous scaffolds (termed MoS) (13). Specifically, a low Dex dosage within the scaffolds decreased the local presence of pro-inflammatory macrophages (termed M1) and promoted an M2-like anti-inflammatory macrophage phenotype, while implants containing elevated Dex dosages inhibited global host cell infiltration and vascularization, resulting in poorly integrated grafts (13). Thus, an optimized design of glucocorticoid release is necessary to balance suppression of inflammation and effective graft integration.

To offset the potentially negative impacts of local Dex release, estradiol (E2), a pro-angiogenic agent that upregulates VEGF gene expression via estrogen receptors α and β, can be co-delivered (14-16). While Dex and E2 could be integrated using standard monolithic dispersion techniques (**Figure 1E**), this approach has limited versatility and typically exhibits early burst release profiles due to the high porosity of the scaffold (17). To address these impediments, we explored a method of encapsulating distinct drug-containing microbeads within high porosity PDMS scaffolds as a dual drug delivery platform (**Figure 1H**). Complementary anti-inflammatory (Dex) and pro-angiogenic (E2) agents were fabricated into discreet drug-eluting PDMS microbeads and integrated into a PDMS scaffold. Drug kinetic profiles of Dex, E2, and Dex+E2 microbeads scaffolds (termed Dex-µBS, E2-µBS, and Dex+E2-µBS, respectively) were characterized, and the impacts of single or dual agent scaffolds on both cytokine secretion from LPS-stimulated macrophages and host cell remodeling within a rodent model were evaluated. Finally, the broader impacts of this drug delivery scaffold system on directing desired host remodeling and integration were discussed.

**Figure 1.**
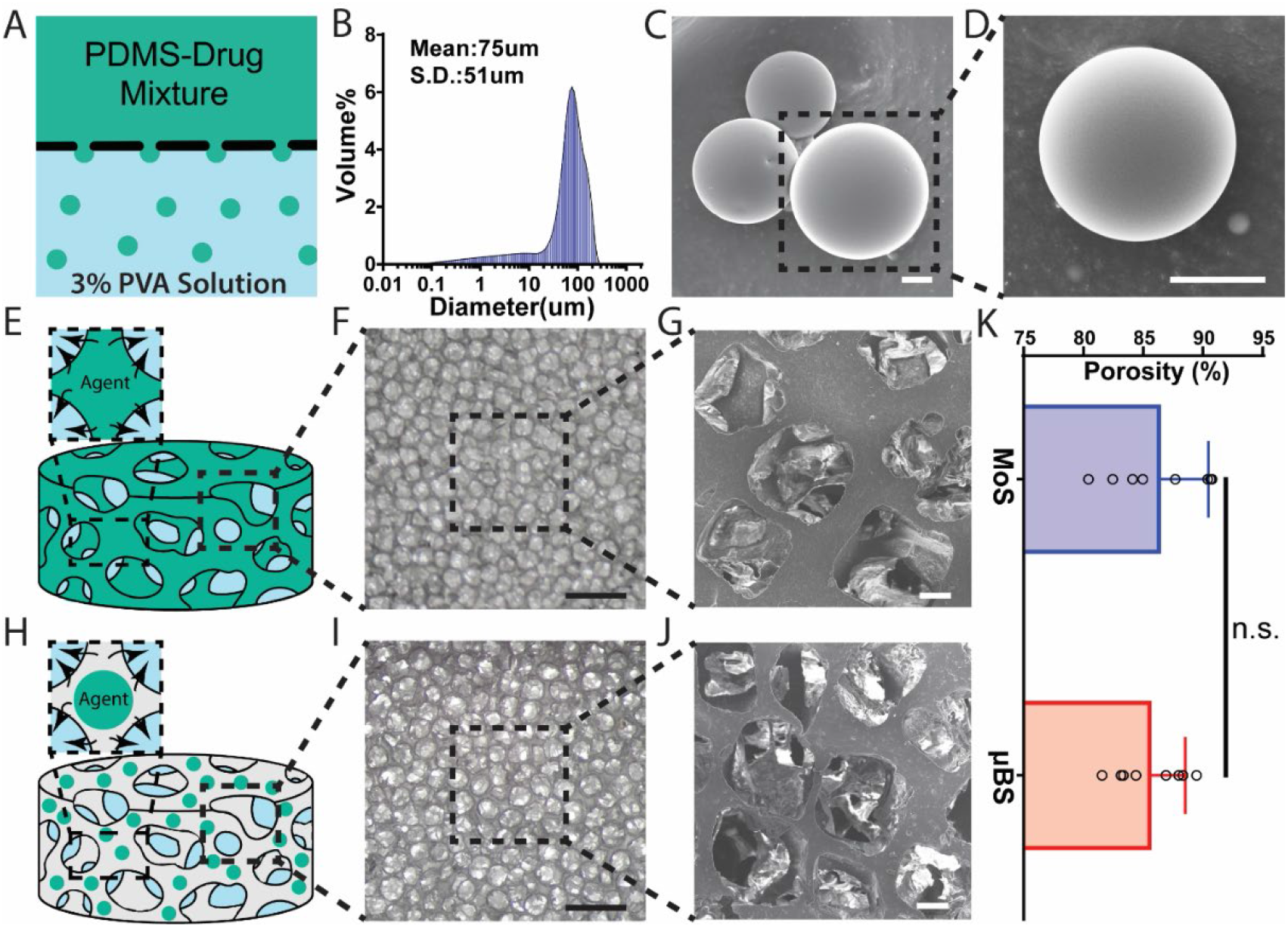
Fabrication and characterization of PDMS scaffolds. **(A)** Schematic of PDMS microbeads fabrication via emulsion method. **(B)** Microbead diameter, as measured by Coulter Counter. **(C-D)** SEM images of PDMS microbeads (*scale bar = 10 μm*). **(E)** Schematic of the monolithic scaffold (MoS). **(F)** Stereomicroscopy image of MoS (*scale bar = 1 mm*). **(G)** SEM images of MoS (*scale bar =100 μm*). **(H)** Schematic of microbead scaffold (µBS). **(I)** Stereomicroscopy image of µBS (*scale bar = 1 mm*). **(J)** SEM image of µBS (*scale bar =100 μm*). **(K)** Scaffold porosity, as measured via mineral oil method (*n = 8 per group; data presented as mean* ± *SD; P* > *0*.*05, not significant, unpaired t test*).

## 2. Materials and Methods

### 2.1 Fabrication and Characterization of PDMS Microbeads

PDMS microbeads were fabricated by an emulsion method. In brief, RTV 615 PDMS (Cat. No. 9480; Momentive) was prepared by mixing Part A and Part B at the ratio of 4:1 in an automatic mixer (Model No. ARE-310; THINKY). Dexamethasone (Cat. No. BML-EI126-0001; Enzo) or 17β-Estradiol (Cat. No. E2758; Sigma Aldrich) were then mixed with the PDMS to form a drug-PDMS slurry. Drug concentration in this slurry was determined by the equation (1):

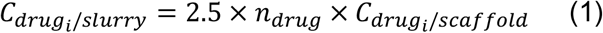

where 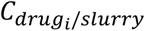 is the drug *i* concentration in the slurry for microbead fabrication, *n*_*drug*_ is the number of drug types in the scaffold, and 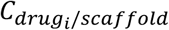 is the desired drug *i* concentration in the scaffold. For instance, the slurry contained 0.125% wt/wt Dex for a Dex microbead scaffold with 0.05% wt/wt Dex or 0.25% wt/wt Dex for a Dex-E2 microbead scaffold with 0.05% wt/wt Dex.

This slurry was then emulsified in a 3% polyvinyl alcohol (PVA; Cat. No. 151937; MP Niomedicals) aqueous solution, as previously reported (18), and filtered through a 41 μm filtration system (Cat. No. SCNY00040; Millipore) to form microdroplets (**Figure 1A**). These suspended microdroplets were then cured in the 3% PVA solution at 80 °C for at least 2 hours. A 100 μm filtration system (Cat. No. SCNY00100; Millipore) was further utilized to select microbeads with a diameter < 100 μm. Microbeads size was characterized by Coulter Counter (Coulter LS13320) and scanning electron microscopy (SEM; Hitachi SU5000 Schottky Field-Emission).

### 2.2 Fabrication and Characterization of PDMS Scaffolds

PDMS scaffolds were fabricated using particle leaching to form scaffolds with an average porosity of 85%, as described previously (2). To incorporate drugs into the scaffold, either a desired amount of drug was directly mixed into PDMS (termed “monolithic scaffolds or MoS”) or drug-containing microbeads were added into PDMS (termed “microbead scaffolds or µBS”) at the ratio determined by the equation (2):

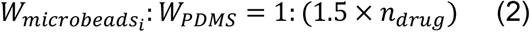

where *W*_*microbeads*_ is the weight of microbeads containing drug *i, W*_*PDMS*_ is the weight of liquid PDMS, and *n*_*drug*_ is the number of the drug types in the scaffold. For instance, Dex-µ BS was fabricated by mixing 1 unit of Dex microbeads into 1.5 units of PDMS, while Dex+E2-µBS was fabricated by mixing 1 unit of Dex and 1 unit of E2 into 3 units of PDMS (so *W*_*total microbeads:*_ *W*_*PDMS*_ would still be 1:1.5) to maintain the same scaffold structure and geometry.

Sodium chloride crystals (Cat. No. s271-500; Fisher Scientific) of 250–425 um were then added to serve as porogen. The resulting paste was then fully mixed and degassed using a THINKY mixer. Completed mixtures were then added to molds (25 mm diameter; 1.5 mm thickness) and cured overnight at 40**°**C at 800 PSI using a compression molder (Desktop Pellet Press; Across International). Disks were then leached in 1.5 L of deionized water for 72 hours, with water refreshed twice daily. After a drying period of 24 hours, 10 mm diameter scaffolds were cut from each disk for the *in vitro* release study. A quarter of the 10 mm diameter scaffold (termed “transplant volume size”) was used for other assessments due to the capacity limit at the transplantation site. Pore size and 3D structures of the scaffolds were examined by SEM and Nano-CT scanning (GE V|TOME|X M 240). The porosity of the scaffolds was tested by a mineral oil method. Briefly, mineral oil (Cat. No. 330779; Sigma Aldrich) of known density was soaked into a piece of a scaffold of known weight and volume, and the porosity was calculated by the equation (3):

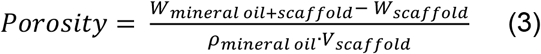

where *W*_*mineral oil*+*scaffold*_ is the total weight of the scaffold soaked with mineral oil, *W*_*scaffold*_ is the weight of the scaffold, *V*_*scaffold*_ is the volume of the scaffold, and *ρ*_*mineral oil*_ is the density of the mineral oil (0.838 g/mL at 25 °C).

### 2.3 Characterization of Drug Release Kinetics *in Vitro*

Dex *in vitro* release kinetics was measured as described previously (13). In brief, a 10 mm disk was immersed in the 1 mL 1% benzalkonium chloride (BKC; B6295; Sigma-Aldrich) solution on a plate shaker (Thermo Fisher) at 300 RPM for 30 days, with daily solution collection and replacement. BKC solution was used to increase the solubility of Dex to avoid saturation during the release period for the higher Dex dosages (24 hr) (13, 19). E2 *in vitro* release kinetics were conducted in the same method but with 1X PBS solution. Release kinetics from dual drug (Dex and E2) scaffolds, as well as the 0.05% wt/wt Dex comparison scaffold, were conducted in 1X PBS (phosphate-buffered saline; Cat. No. 10010023; Gibco). Dex from the high Dex loading scaffolds (1% and 0.5% wt/wt) was measured by high-performance liquid chromatography (HPLC; Thermo Fisher) with an Acclaim 120 C18 3 µm 4.6×150 mm column ((Cat. No. DX059133; Thermo Scientific). The mobile phase (50% acetonitrile and 50% water) was run at 1.0 ml/min for 9 min with a UV detection at 240 nm, as modified from other protocols (20). Three independent replicates were procured from a single injection for each scaffold group. Dex from lower Dex loaded scaffolds (0.25%, 0.1%, and 0.05% wt/wt) was measured by a Dex enzyme-linked immunosorbent assay (ELISA; Cat. No. 101516; Neogen) kit. E2 was measured by an E2 ELISA kit (Cat. No. ADI-901-008; Enzo). N = 3 independent replicates were conducted for each scaffold group with an n = 2 technical replicates run per sample for each drug measurement. Release results were presented as ng per day per transplant volume size scaffold.

Cumulative results were calculated by stacking daily release values, while the release values between two nonconsecutive collection points were estimated by the average of these two data points. Zero-order and first-order release models were used to analyze the data using the equations (4) and (5) in GraphPad Prism 8 software:

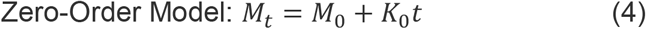

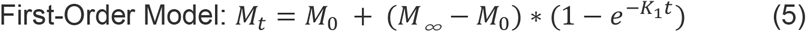

where *M*_*t*_ (ng) is the cumulative amount of drug released at time t (day), *M*_0_ (ng) is the initial amount of drug in the solution, *M*_∞_ (ng) is the total amount of drug that could be released from the scaffolds, *K*_0_ (ng/day) is the zero-order release rate, and *K*_1_ (1/day) is the first-order release rate (21). Additional modeling used to predict targeted E2 dosages is summarized in Supplementary Text 1 (**Table S2**).

### 2.4 Isolation and Differentiation of Bone-marrow Derived Macrophages

Bone-marrow derived macrophages were isolated from 8 to 12-week-old C57BL/6 mice (Jackson Laboratory) under protocols approved by the Institutional Animal Care and Use Committee (IACUC) at the University of Florida. Following euthanization, bones were harvested from the limb site and cleaned by 0.75% vol/vol chlorohexidine. The marrow was flushed out of the bones via 2 - 5 mL 4 °C DPBS (Dulbecco’s Phosphate-Buffered Saline; Cat. No. 21-031-CV; Gibco) injection. The resulting pellet was filtered through a 70 µm cell culture strainer (Cat. No. 352350; Corning) and centrifuged at 500 x g for 10 minutes. Cells (4 × 10^6^) were seeded on 80 mm non-tissue culture treated plates with 10 mL of DMEM/F-12 media (Cat. No. 10565018; Gibco) which contains 100U/mL M-CSF (Cat. No. 315-02; Peprotech). On day 3 of culture, an additional 5 mL of media was added to each dish to continue the differentiation process. After 7 days, bone-marrow derived macrophages were detached from the plates by using CellStripper solution (Cat. No. 25056CI; Corning) and collected in the cold DMEM/F-12 media. Cell scrapers were used to help this process. The cell suspension was then centrifuged at 400 x g for 10 minutes at 4 °C to remove any CellStripper residues, and the harvested cells were cultured for the following experiments.

### 2.5 Macrophages’ Inflammatory Responses to Scaffolds *in Vitro*

Following one-week culture, bone-marrow derived macrophages were seeded at 100,000 cells per 3.5 cm^2^ in a tissue-treated 6 well plate and incubated in 500 uL DMEM/F-12 media containing 10 ng/mL lipopolysaccharide (LPS) (Cat. No. L4391; Sigma Aldrich) to skew the cells towards a pro-inflammatory M1 macrophage phenotype, as previously described (22). Scaffolds (transplant volume size) were also added to macrophage cultures (N = 4 per treatment group). Scaffold groups included: drug-free control scaffolds, 0.05% wt/wt Dex-µBS, 0.01% wt/wt E2-µBS, and combination scaffolds (0.05% wt/wt Dex + 0.01% wt/wt E2-µBS). Following 24 hours of culture at 37°C, media was collected and analyzed using a MILLIPLEX® Mouse Cytokine Magnetic Bead Panels (EMD Millipore) kit for characterization of cytokine release, i.e. IL-6, IL-10, VEGF, and TNF-α, following the manufacturer instructions. Drug-free control and quality control groups were also appropriately included. N = 4 independent replicates were conducted for each scaffold group with an n = 2 technical replicates run per sample for each group. Data acquisition was performed with Luminex MAGPIX® instrument on xPONENT® 4.2 software platform. Quantification of cytokine levels was done with MILLIPLEX® Analyst 5.1 software. Results were pooled from two independent studies.

### 2.6 Biocompatibility Transplantation within the Epididymal Fat Pad Site

All biocompatibility implants used 12-week-old male C57BL/6J mice (Jackson Laboratory) under protocols approved by IACUC at the University of Florida. Drug scaffolds (one-quarter of a 10 mm disk) were sterilized in 70% isopropanol for 30 minutes, followed by 5 minutes 1X PBS wash for 5 times, as conducted previously (13). Scaffolds were then incubated with 250 μg/mL human plasma fibronectin (Cat. No. 33016-015; Gibco) overnight at 37 °C to enhance hydrophilicity, as previously described (2, 13). Following 3 independent washes in 1X PBS, scaffolds were placed within mobilized epididymal fat pads (EFP), wrapped in the fat pad tissue, and sealed with fibrin gel (fibrinogen at 7.5 mg/mL, aprotinin at 170 μg/mL, thrombin at 10 U/mL, CaCl_2_ at 9mM, and HEPES at 0.36mM), as previously described. Scaffold groups of different structures (monolithic and microbead) with different Dex concentrations (1%, 0.5%, 0.25%, 0.1%, and 0% wt/wt) were used to screen the microbead scaffold prototype in a 14-day study (N = 1 per group). Microbead scaffold groups of different drugs (drug-free, 0.05% wt/wt Dex, 0.01% wt/wt E2, 0.1% wt/wt E2, and combination scaffolds (0.05% wt/wt Dex + 0.01% wt/wt E2)) were included in a 10-day study to examine the dual drug release effect (N = 3 per group). Animals were monitored daily until the termination of the study. For vascularization studies, DyLight ® 594 lectin (Cat. No. DL-1177; Vector Laboratories) was given to the animals at the dose of 2 μg/g via intravenous (IV) injection, and the grafts were retrieved after 30 minutes and fixed in 10% formalin solution (Cat. No. SF98-4; Fisher Chemical) before whole-mount imaging. All other retrieved grafts were fixed in 10% formalin solution and further histologically processed and paraffin-embedded before being sliced. Histological staining included Masson trichrome stain (Cat. No. 87019; Richard-Allan Scientific) and α-smooth muscle actin (α-SMA) (primary Rabbit anti-mouse alpha-SMA antibody; Cat. No. ab5694; Abcam; 1:50 dilution; secondary Goat anti-rabbit antibody; Cat. No. A11008; Life Technologies; 1:150 dilution). Fluorescent images were acquired using a Leica SP8 confocal microscope. Terminal blood collection was conducted via retro-orbital bleeding from animals under anesthesia with 3.2% wt/vol sodium citrate (Cat. No. S4641; Sigma Aldrich) anticoagulant solution. The blood samples were immediately centrifuged at 1000 rpm for 10 min after collection to obtain plasma and stored at 4 °C prior analyses. Dex ELISA kit (Cat. No. 101516; Neogen) and E2 High sensitivity ELISA kit (Cat. No. ADI-900-174; Enzo) were used to detect drug concentration in N = 3 independent blood samples for each scaffold group with an n = 2 technical replicates run per sample.

### 2.7 Image Analysis

NanoCT images (n = 6 for each group) were taken at 6 different time points (t = 1, 3, 5, 7, 9, 11 s) from the videos (**Supplemental Video S1-2**) and processed in Fiji ImageJ software to calculate the porosity from the binary area. Nuclei counts from histology images (N = 1 for the 14-day experiment; N = 2 for 0.1% E2 group; N = 3 for all other groups; n ≥ 5 images for each sample) were detected by a customized Python 2.7 code developed in Jupyter Notebook in Anaconda. Briefly, histology images were turned into binary images with the threshold adjusted to only show nuclei. A Gaussian filter was applied to smooth the images before regional maxima were found to estimate the nuclei counts (as outlined http://pythonvision.org/basic-tutorial/). The intensity of staining in trichrome images was analyzed in Fiji ImageJ software by turning the images into the binary and adjusting threshold to remove the scaffold background. The intensity and area of α-SMA staining (N = 1 for each group; n ≥ 4 images for each sample) were processed in Fiji ImageJ software as well. Perfused lectin images (N = 1 for each group; n ≥ 3 images for each sample) were analyzed in AngioTool software with a setting of “Vessel Thickness = 15”, “Small Particles = 800”, and “Fill Holes = 0”.

### 2.8 Statistical Analysis

In all experiments, the results are expressed as mean ± SD. When only two groups were compared, an unpaired t test was performed, as designed. For more than two groups comparisons, one-way ANOVA was conducted with Tukey’s post hoc. For kinetic release studies, two-way RM ANOVA (time and drug dosage as factors) was conducted with post hoc Sidak’s multiple comparisons test. For mathematical modeling results and Dex-only 14-day explant histology analysis, two-way ANOVA (scaffold format and drug concentration as factors) was conducted with post hoc Tukey’s multiple comparison test. For all statistical analyses, a P < 0.05 was considered significantly different. All statistical analyses were conducted in GraphPad Prism 8 software. The details of the statistical results can be found in **Table S3**.

## 3. Results

### 3.1 Characterization of PDMS microbeads and Scaffolds

For the controlled release of multiple drugs, a PDMS-based macroporous scaffold platform with embedded microbeads was developed. To retain the structure of the scaffold, PDMS microbeads were fabricated using an emulsion technique (**Fig 1A)**; resulting PDMS-based microbeads exhibited an average size of 75 ± 51 µm, with SEM imaging confirming their spherical nature (**Figure 1B-D)**. The integration of 40% wt/wt microbead within the macroporous scaffold had no impact on scaffold structure or porosity when compared to monolithic scaffolds. Stereomicroscopy and SEM images illustrated a highly porous scaffold with pore size and interconnectivity visually comparable to control scaffolds (**Figure 1F-G & I-J**) (2, 13). Global scaffold porosity of the microbead scaffolds (85.6 ± 2.9%) was statistically identical to that of the control monolithic scaffolds (**Figure 1K**). NanoCT scanning (**Figure S1** and **Supplemental Video S1-2**) further confirmed similar interconnectivity in 3D and comparable global porosity (**Figure S1E**).

### 3.2 Characterization of Dex Releases from PDMS Microbeads Scaffolds

The impact of the microbead scaffold approach on drug kinetics was evaluated by comparing Dex release from either a monolithic (MoS) or microbead scaffold (µBS) platform at four distinct drug doses (1%, 0.5%, 0.25%, and 0.1% wt/wt) (**Figure 2B-C**). The drug release curves of Dex-MoS were significantly distinct from Dex-µBS curves at the loadings of 1%, 0.5%, and 0.25% wt/wt (P = 0.0012, P = 0.0037, P = 0.0004, respectively) over the 30-day collection period, while the 0.1% wt/wt Dex loading was only slightly insignificant (P = 0.053). A zero-order release model was employed to classify initial burst release trends (days 0-5). This model had a goodness-of-fit R^2^ value > 0.89 for all the fitting curves except for the 0.1% wt/wt MoS group (R^2^ = 0.76; detailed in **Table S1**). As expected, scaffold format and drug concentration, as well as their interaction, imparted significant impacts on the initial burst release (P < 0.0001 for all three factors), as measured by the zero-order release rate constant *K*_0_. Furthermore, at Dex drug loadings of 1% and 0.5% wt/wt, monolithic scaffolds (MoS) exhibited a significantly higher burst release rate than their microbead scaffold (μBS) counterparts (**Figure 2D**). The durability of drug release from the two platforms for the four dosages was characterized by employing a first-order release model for the full 30-day release window; the model had a goodness-of-fit R^2^ value > 0.85 for all groups and doses except for the 0.1% wt/wt MoS group (R^2^ = 0.71; **Table S1**). Based on these models, scaffold format had a major impact on the overall first-order release rate (P < 0.0001), while drug concentration and their interaction contributed less influence (P = 0.029 and 0.013, respectively). For the same scaffold format (MoS or µBS), the Dex drug concentration imparted no significant difference, indicating that this range of drug doses did not alter the durability of drug release. The MoS platform exhibited a significantly higher first-order release rate, *K*_1_, than its corresponding Dex-µBS for loadings of 0.5% and 0.25% wt/wt (**Figure 2G**). Globally, the µBS format reduced the first-order release rates in a manner that prolonged half-max times by at least 1.6-fold than corresponding MoS (**Figure 2H&I** and **Table S1**). When the Dex loading in the µBS platform was further decreased from 0.1% to 0.05% wt/wt (**Figure 2C**), the zero-order burst release rate was significantly reduced (P < 0.0001; **Figure 2D**), while the first order long-term release rate remained similar (P = 0.64; **Figure 2G**). Overall, the µBS platform dampened burst release and extended the durability of drug release when compared to the standard MoS platform.

**Figure 2.**
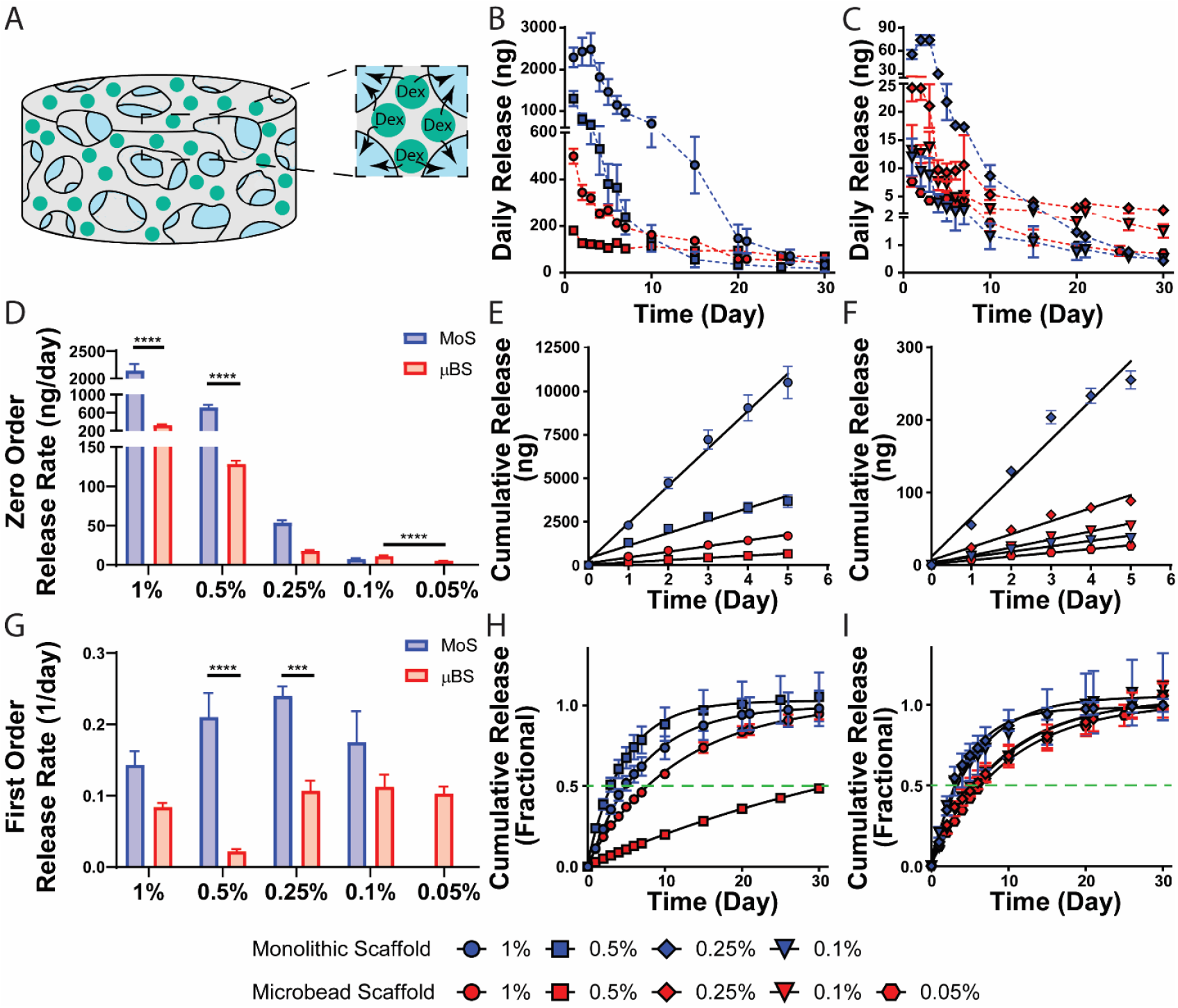
Drug release kinetics and analyses of Dex-releasing scaffolds. **(A)** Schematic of Dex-µBS; **(B-C)** Summary of daily Dex release *in vitro* for MoS (blue) or µBS (red) formats containing 1 and 0.5% **(B)** and 0.25, 0.1, and 0.05% **(C)** wt/wt Dex (*n = 3 samples for each group; data presented as mean* ± SD; *dashed lines – the connection between data points to illustrate trends*). **(D-F)** Zero-order release analyses for the first five days: **D)** Summary of release rate K_0_ values derived from individual release curves; Cumulative release curves for high **(E)** and low **(F)** Dex drug loadings with lines representing zero-order model predictions and points illustrating experimental data set (*n = 18 data points for each group; data presented as mean* ± *SEM*) **(G-I)** First-order release analyses for the 30-day release period: **G)** Summary of release rate K_1_ values derived from individual release curves; Cumulative release curves for high **(H)** and low **(I)** Dex drug loadings, normalized to predicted M_∞_ values, with lines representing first-order model predictions and points illustrating experimental data set (*n = 39 - 42 data points per group; data presented as mean* ± *SEM*). *Two-way ANOVA with Tukey’s post-hoc*, \*\*\**P* < *0*.*001*, \*\*\*\**P* < *0*.*0001*

### 3.3 Evaluation of Dex Monolithic and Microbeads Scaffolds *in Vivo*

To characterize the impact of these distinct Dex release profiles on the host response, scaffolds were implanted into the epididymal fat pad of C57BL/6J mice. Explants were collected after 14 days and trichrome stained to capture cellular migration and extracellular matrix deposition, with images collected at both the host tissue-scaffold interface and within the central region of the implant (**Figure 3**). Overall, scaffold type, drug concentration, and their interaction had significant impacts on cellular infiltration into the center of the scaffolds (P < 0.0001 for all factors). At the highest Dex loadings (1% and 0.5% wt/wt), both the MoS and µBS platforms globally reduced cellular infiltration into the center of the scaffold, when compared to drug-free controls, with no significant differences between the scaffold format (**Figure 3Q**). Decreasing the Dex loading from 0.5 to 0.25% wt/wt resulted in elevated host cell presence within the MoS (P = 0.007). Contrarily, decreasing drug dosage from 0.5 to 0.25% for the µBS format did not alter the suppressive features of the implant (P = 0.90). At the 0.1% wt/wt Dex loading, MoS implants did not significantly impact host cell migration when compared to the drug-free control group (P = 0.83), while explants from 0.1% wt/wt µBS platform still significantly hindered host cell infiltration (P < 0.0001). Overall, µBS implants decreased host cell infiltration in a dose-dependent manner (P = 0.0002; one-way ANOVA). Reducing the drug loading of the µBS further to 0.05% still imparted significant suppression of host cell infiltration when compared to drug-free controls (**Figure 4**), although explants from this study were collected at an earlier timepoint (10 days post-transplant).

**Figure 3.**
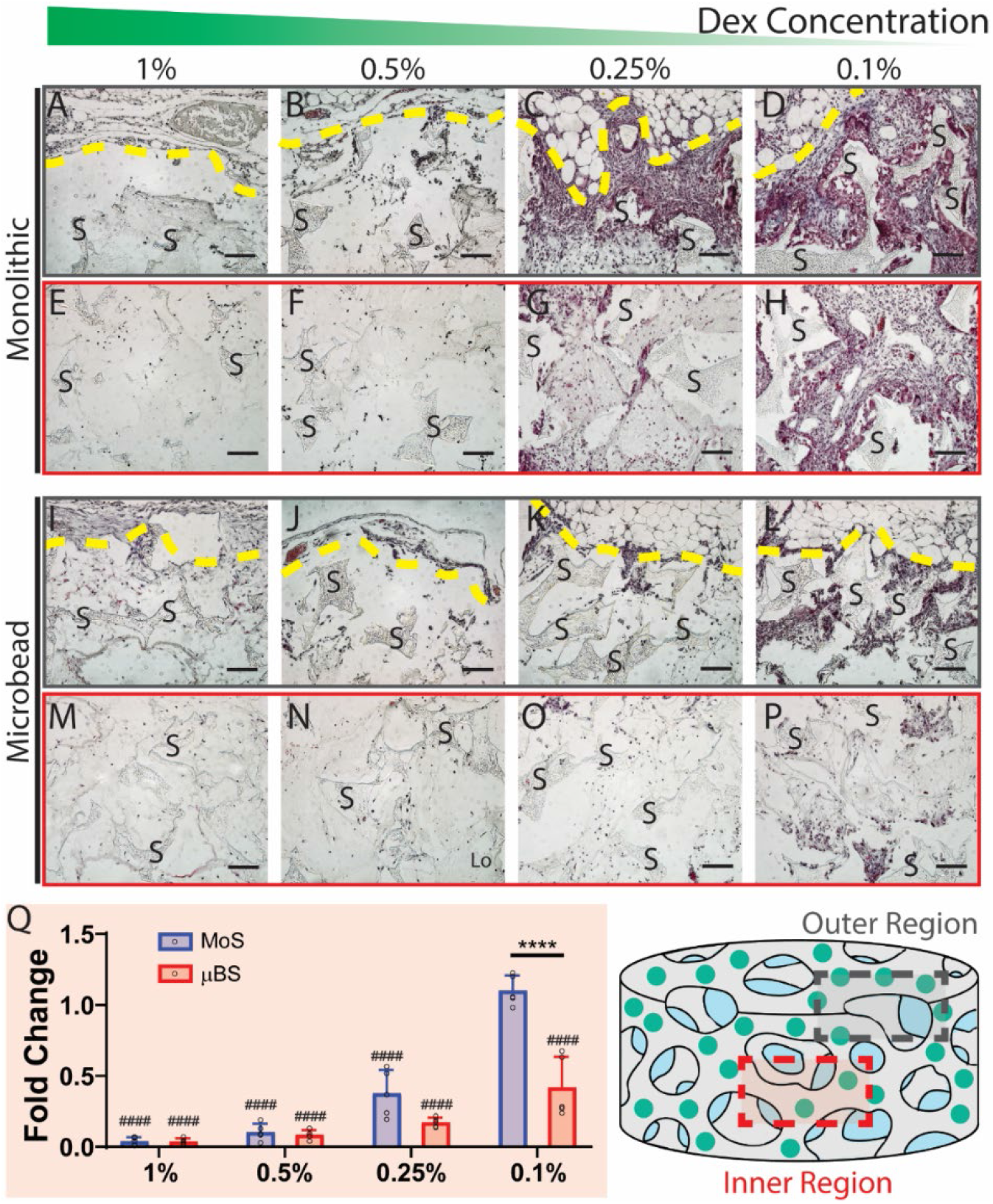
Trichrome staining of Dex-loaded scaffolds explanted from epididymal fat pad 14 days post-implantation. Trichrome stained cross-section of 1%, 0.5%, 0.25%, and 0.1% (wt/wt) Dex-MoS collected from the host/scaffold interface (**A-D;** designated by schematic as the “Outer Region”) or the center of the scaffold implant (**E-H**; designated by schematic as the “Inner Region”) *Yellow dashed line = scaffold/host interface*. Trichrome stained cross-section of 1%, 0.5%, 0.25%, and 0.1% (wt/wt) Dex-µBS collected from the host/scaffold interface (**I-L;** designated by schematic as the “Outer Region”) or the center of the scaffold implant (**M-P**; designated by schematic as the “Inner Region”) *Yellow dashed line = scaffold/host interface; S = scaffolds; scale bar = 100 μm*. **(Q)** Fold change in nuclei counts, as quantified from processed images of the Inner Region and normalized to the 0% control group (*n = 5 for each group; data presented as mean* ± *SD;* \*\*\*\**P* < *0*.*0001, two-way ANOVA with Tukey’s post-hoc; one-way ANOVA with Tukey’s post-hoc*, ^*####*^*P* < *0*.*0001 compared to drug-free control*).

**Figure 4.**
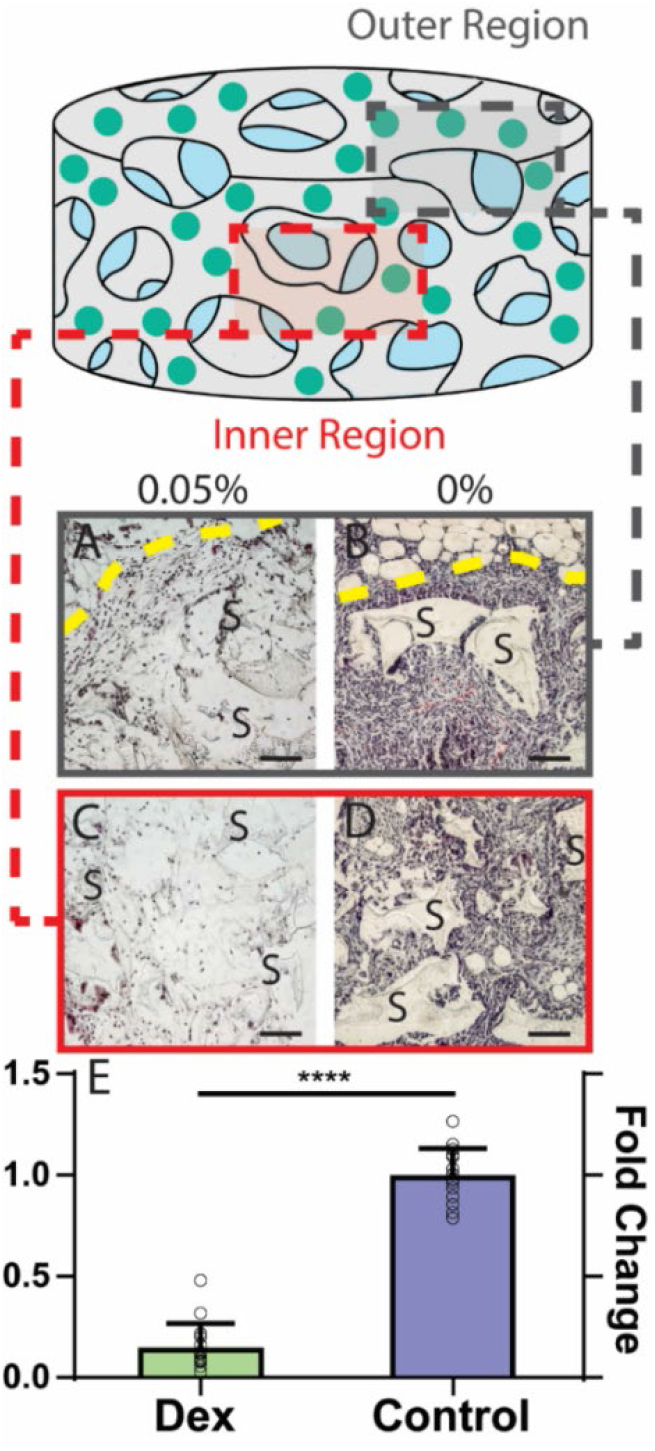
Trichrome staining of 0.05% wt/wt Dex and drug-free microbead scaffolds explanted from epididymal fat pad 10 days post-implantation. **(A-B)** Trichrome stained cross-section of 0.05 and 0% (wt/wt) Dex-µBS collected from the host/scaffold interface (**A-B;** designated by schematic as the “Outer Region”) or the center of the scaffold implant (**C-D**; designated by schematic as the “Inner Region”) *Yellow dashed line = scaffold/host interface; S = scaffolds; scale bar = 100 μm*. **(E)** Fold change in nuclei counts, as quantified from processed images of the Inner Region and normalized to the 0% control group (*n = 15 for each group; data presented as mean* ± *SD;* \*\*\*\**P* < *0*.*0001, unpaired t test*).

### 3.4 Evaluation of E2 Microbeads Scaffolds *in Vitro* and *in Vivo*

To explore the potential of the microbead platform to serve as a controlled release system for other agents, the pro-angiogenic agent estrogen (E2) was formulated into PDMS microbeads and the controlled release from the µBS format was characterized. Two concentrations of E2 were selected (0.01% and 0.1% wt/wt E2). The lower dosage was based on generated µBS predictive models and pro-angiogenic doses sourced from published reports (16, 23) (for further details see **Supplementary Text 1**). As shown in **Figure 5A**, durable E2 release was observed over the course of 30 days, ranging from 3.3 – 35.4 ng/day for the 0.1% wt/wt E2-µBS and 0.05 – 8.1 ng/day for the 0.01% wt/wt E2-µBS. To characterize the impact of local E2 release on host responses to the scaffold implant, another biocompatibility implant study was conducted, with explant analysis performed 10 days post-implantation to capture early-stage angiogenesis. As shown in **Figure 5B-D**, trichrome stained E2-µBS implants exhibited altered host responses. Of interest, the lower dosage E2-µBS (0.01% wt/wt) displayed elevated matrix deposition but not cellular infiltration, compared to drug-free control scaffolds. The higher drug loading of 0.1% E2-µBS, however, decreased cellular infiltration and matrix deposition. Based on these results, the 0.01% E2-µBS was used in subsequent dual-drug release experiments.

**Figure 5.**
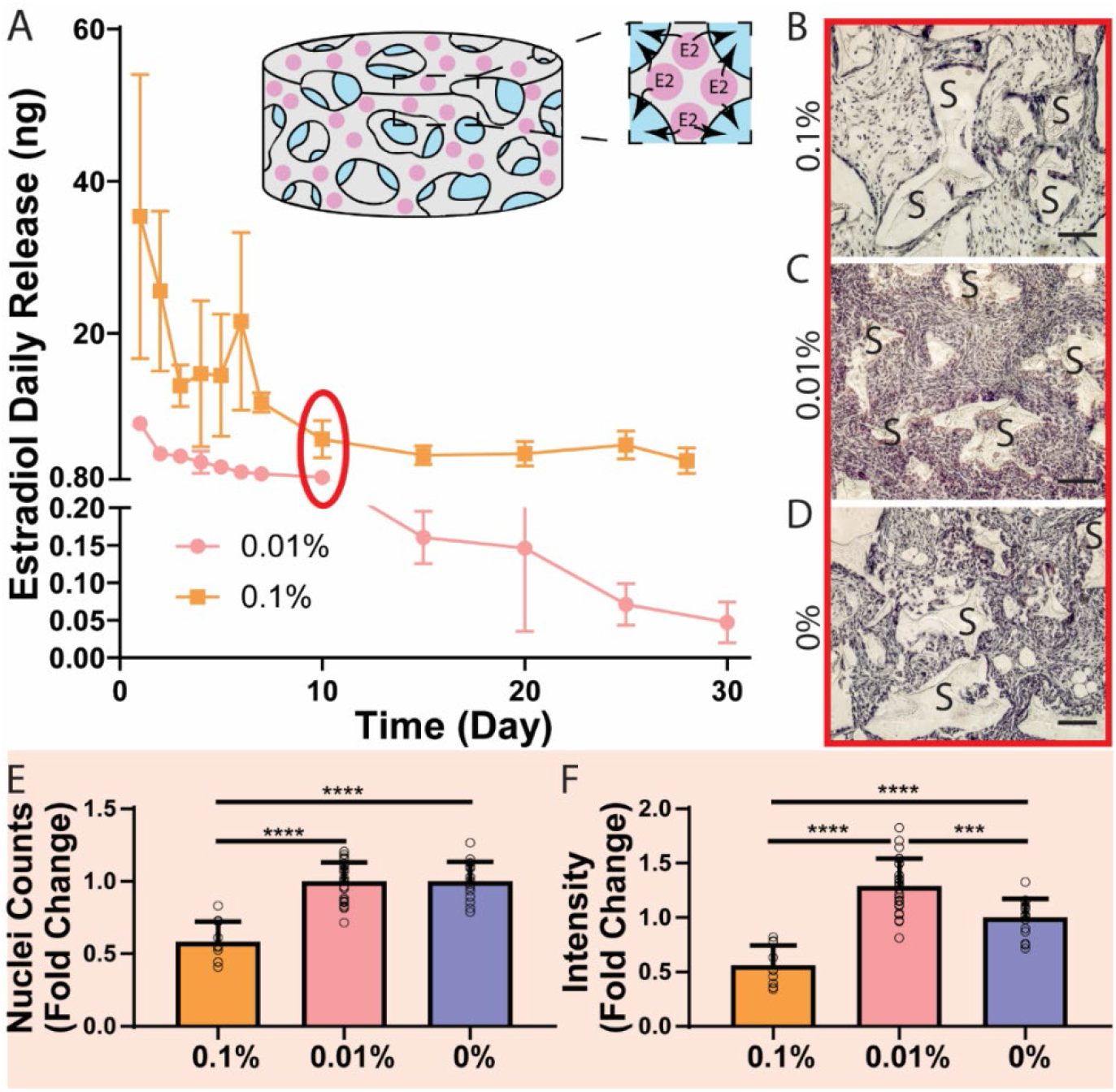
Characterization of estrogen (E2) releasing microbead scaffolds. **(A)** 30-day *in vitro* release profiles for 0.01 and 0.1% E2-μBS (n = 3 for each group). **(B-D)** Trichrome stained images of the center of 0.1%, 0.01%, and 0% (wt/wt) E2-μBS explanted 10 days post-implantation (*S refers to scaffolds; scale bar = 100 μm*). **(E)** Fold change in nuclei counts, as quantified from processed images of the Inner Region and normalized to the 0% control group (*n* ≥ *10 for each group; data presented as mean* ± *SD*). **(F)** Fold change in Trichrome stain intensity, normalized to the 0% control group (*n* ≥ *10 for each group; data presented as mean* ± *SD*). O*ne-way ANOVA with Tukey’s post-hoc* \*\*\**P* < *0*.*001*, \*\*\*\**P* < *0*.*0001*

### 3.5 Characterization of Dual Drug Release from PDMS Microbeads Scaffolds

Following the kinetic evaluation of single-drug microbead scaffolds, E2 and Dex microbeads were then co-integrated into a single scaffold and their subsequent kinetic release curves were assessed. The low dosages of Dex (0.05%) and E2 (0.01%) were selected due to their inferred combinatorial effects. To avoid inducing an changes in the structure of the macroporous scaffold, the amount of Dex or E2 drug per PDMS bead was doubled, which permitted stability in the total number of beads loaded per scaffold with an equal number of Dex and E2 microbeads. The overall kinetic release curves between the single agent and double agents were equivalent for each drug (P = 0.16 and 0.052 for Dex and E2, respectively; **Figure 6B&C**). Modeling the kinetic release, the zero-order model showed significantly reduced release rate constant (K_0_), when compared to single agent µBS (**Figure 6D&E**); the first-order release rate of Dex was also reduced by the combination (**Figure 6F**).

**Figure 6.**
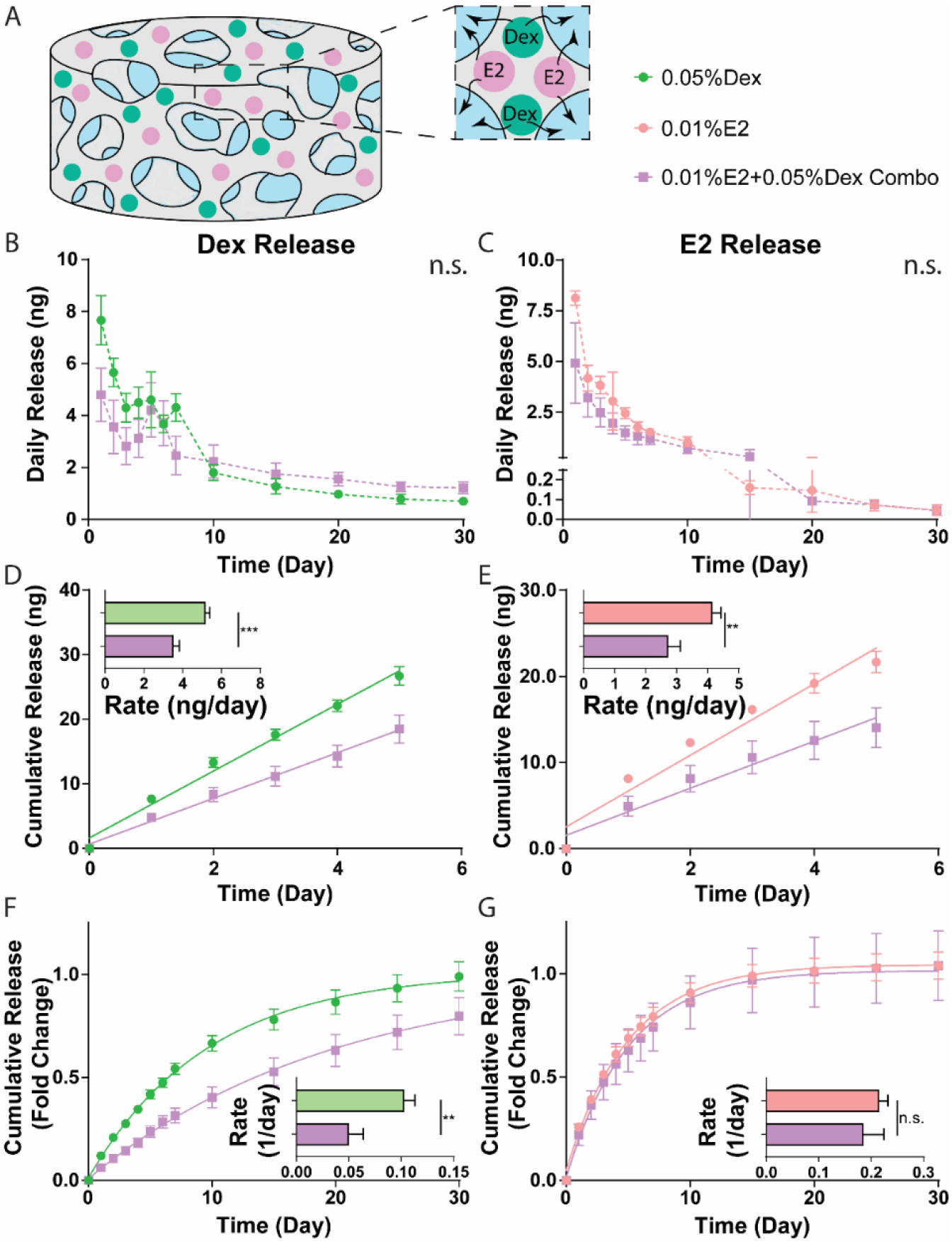
Characterization of dual drug release of dexamethasone and estrogen from microbead scaffolds. **(A)** Schematic of the combination scaffold. **(B-C)** 30-day *in vitro* Dex **(B)** and E2 **(C)** release profiles (*n = 3 for each group; two-way ANOVA*). **(D-E)** Zero-order release analyses for the first five days: Cumulative release curves for Dex **(D)** and E2 **(E)** with lines representing zero-order model predictions and points illustrating experimental data set (*n = 18 data points for each group; data presented as mean* ± *SEM*); Insets: Summary of zero-order release rate K_0_ values derived from individual release curves for Dex **(D)** and E2 **(E). (F-G)** First-order release analyses for the 30-day release period: Cumulative release curves for Dex **(F)** and E2 **(G)**, normalized to predicted M_∞_ values, with lines representing first-order model predictions and points illustrating experimental data set; Insets: Summary of first-order release rate K_1_ values derived from individual release curves for Dex **(F)** and E2 **(G)**. (*n = 39 data points for each group; data presented as mean* ± *SEM*). *D-G statistics: Unpaired t test, n*.*s. = not significant;* \*\**P* < *0*.*01;* \*\*\**P* < *0*.*001*

### 3.6 Modulation of Macrophage Response to Inflammatory Stimuli *in Vitro*

To investigate the combinatorial effects of Dex and E2 on host responses, murine bone marrow-derived macrophages were co-incubated with single agent, double agent, or drug-free scaffolds (0.05% Dex-µBS; 0.01% E2-µBS; 0.05% Dex + 0.01% E2-µBS; no drug scaffold). Macrophages were concurrently treated with LPS to induce a pro-inflammatory M1 phenotype (22, 24). After 24 h *in vitro*, cell culture media was characterized for cytokine content, with a focus on classic pro-inflammatory (IL-6 and TNF-α) and pro-healing (IL-10 and VEGF) cytokines. As summarized in **Figure 7**, the 0.05% Dex-µBS had the most potent and pleiotropic suppressive effect, with significant dampening of all pro-inflammatory and pro-healing cytokines tested. The E2-µBS platform had no significant impact on cytokine secretion, when compared to drug-free control scaffolds. When Dex and E2 were combined, however, the inclusion of estrogen dampened Dex suppression of pro-healing cytokines with IL-10 levels comparable to control scaffolds and VEGF levels less suppressed than Dex only levels. Results indicate that Dex has a dominant effect on LPS-induced M1 macrophages *in vitro*, but the co-delivery of E2 can restore selected pro-healing macrophage features.

**Figure 7.**
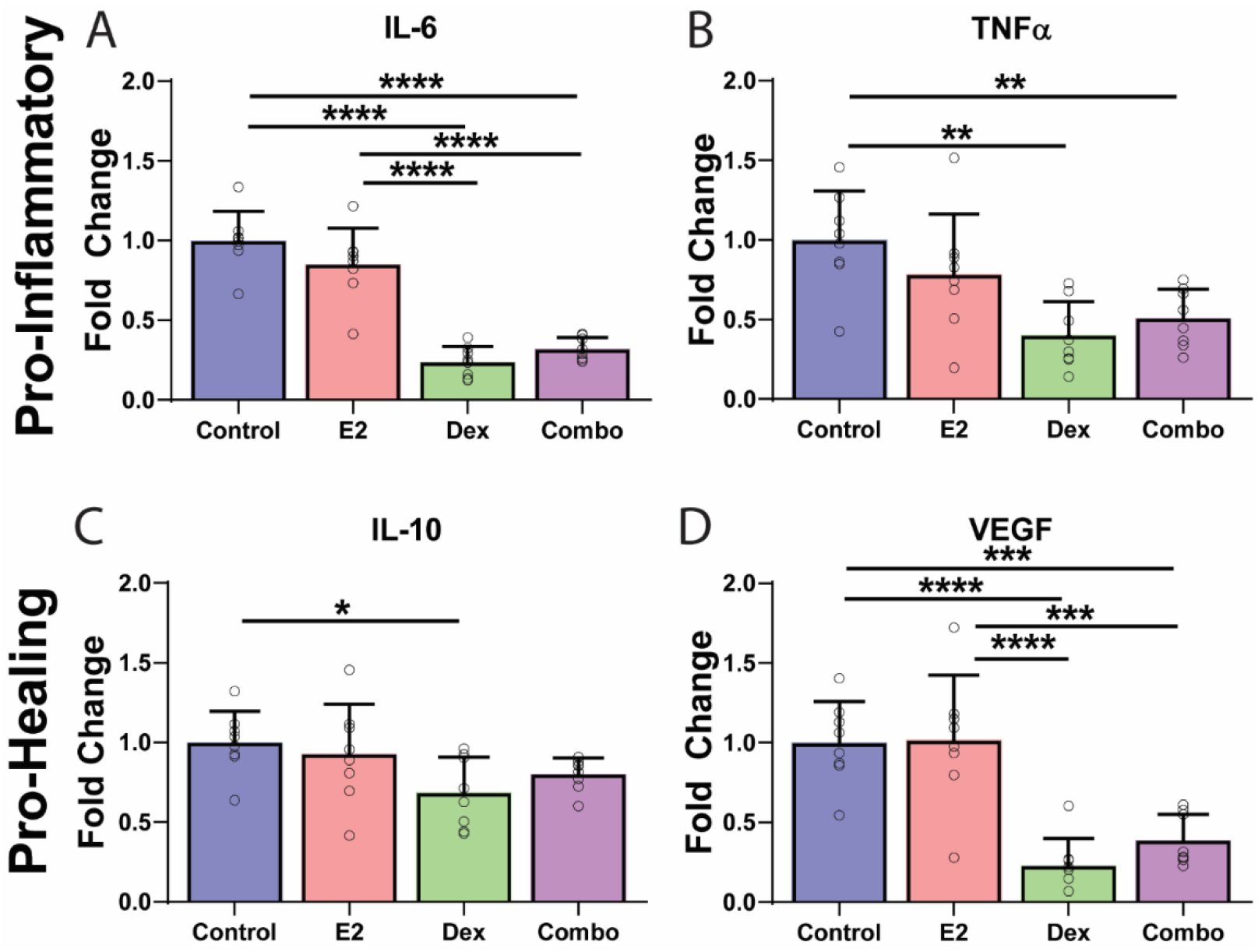
Cytokine analysis of bone-marrow derived macrophages co-cultured without or with drug eluting microbead scaffolds. (0.01% E2 and/or 0.05% Dex). **(A-B)** Fold change in proinflammatory cytokines IL-6 and TNF-alpha, normalized to drug-free μBS. **(C-D)** Fold change in pro-healing cytokines IL-10 and VEGF, normalized to drug-free μBS. (*n = 8 for each group; data presented as mean* ± *SD;* \**P* < *0*.*05*, \*\**P* < *0*.*01*, \*\*\**P* < *0*.*001*, \*\*\*\**P* < *0*.*0001, one-way ANOVA with Tukey’s post-hoc*).

### 3.7 Evaluation of Dual-Drug Delivery Scaffolds *in Vivo*

To examine the impact of this combinatory approach on host responses *in vivo*, scaffolds (drug-free control, 0.01% E2-µBS, 0.05% Dex-µBS, and 0.01% E2 + 0.05% Dex-µBS) were implanted into the EFP site of a C57BL/6 murine model. Grafts were explanted 10 days post-transplant and histologically characterized via trichrome and α-SMA. Competent vascularization was also evaluated via perfused lectin staining. As summarized in **Figure 8** and **Figure S3**, significant alterations in host responses were observed in the center of these implanted scaffolds. Trichrome staining indicated robust host cell infiltration, deposition, and vascularization for both control and E2-µBS implants (**Figure 8A-B**). Alternatively, local Dex release within µBS implants profoundly suppressed host integration, with minimal cellular presence and extracellular matrix deposition (**Figure 8C&E**). The dual drug release implants, however, illustrated a more modest suppression, with evidence of elevated cellular integration and matrix deposition when compared to Dex-only implants (**Figure 8D&E**). This trend was further supported by α-SMA stained sections, with similar suppression in α-SMA staining for Dex-µBS (**Figure 8H&E**). The inclusion of E2, either alone or co-delivered with Dex exhibited opposite effects, with elevated α-SMA when compared to drug-free control implants (**Figure 8F-J**). Competent vascularization was also impacted in a similar manner. While control and E2-µBS implants illustrated robust vascularization, local Dex release suppressed their presence (**Figure 8M&O**). The co-delivery of E2 and Dex, however, resulted in vascularization statistically equivalent to drug-free controls (**Figure 8K-O**). Additional quantifications of vessel characteristics further highlight the benefits of local E2 delivery (**Figure S3**). To assess potential systemic effects of local drug release, plasma Dex and E2 levels were collected at the time of graft retrieval. Regardless of the drug combination, no significant alterations in circulating Dex or E2 levels were measured (**Figure S4**).

**Figure 8.**
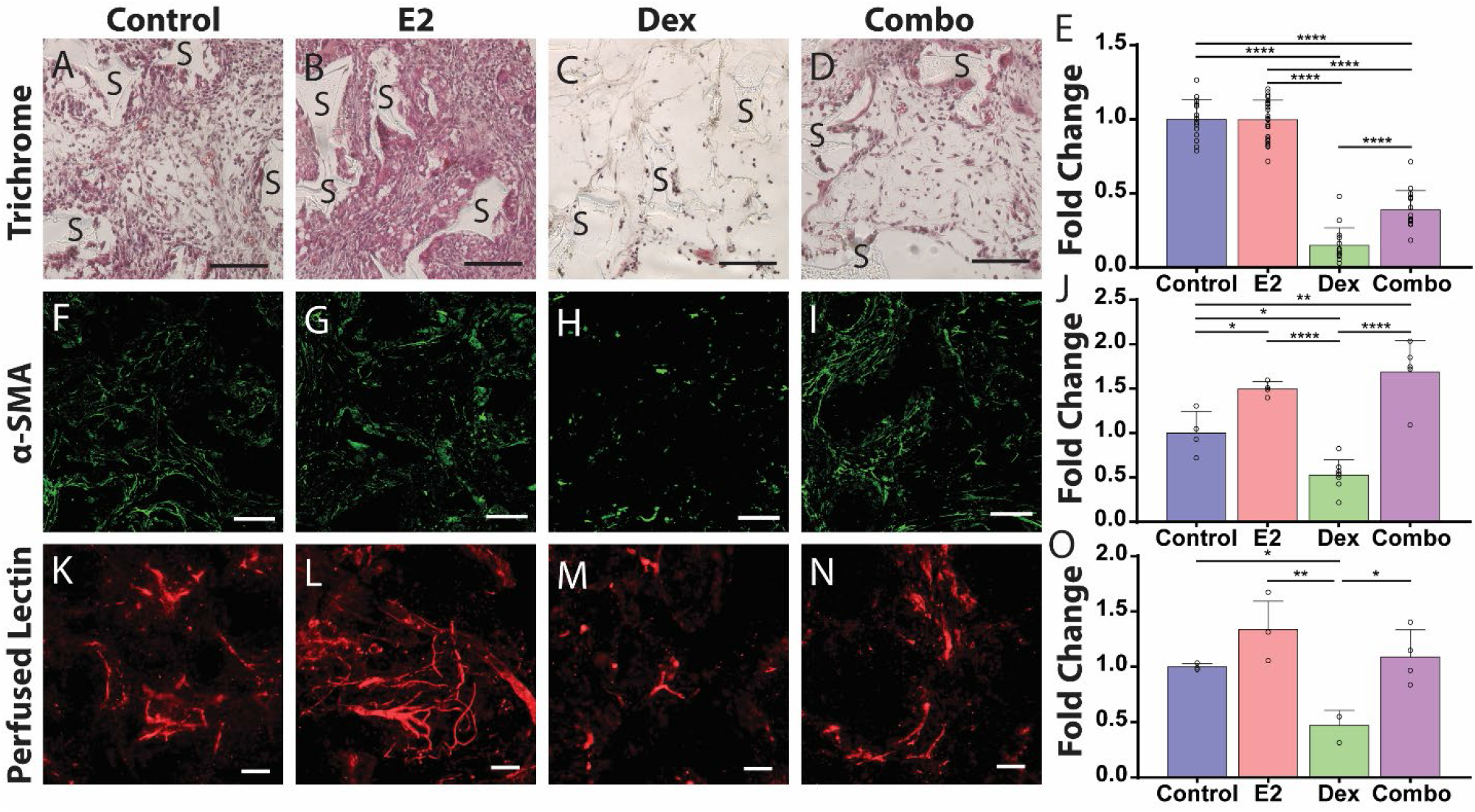
Comparison of host responses to drug-free or Dex and/or E2 eluting microbead scaffolds explanted 10 days post-implantation. (0.05% Dex and/or 0.01%E2). Trichrome **(A-D)**, α-SMA **(F-I)**, and perfused lectin **(K-N)** stained images of the center of drug-free, 0.01% E2, 0.05% Dex, or 0.01% E2 + 0.05% Dex (Combo) μBS (*S refers to scaffolds; scale bar = 100 μm*). **(E)** Fold change in nuclei counts, as quantified from processed images of the Inner Region and normalized to the 0% control group (*n* ≥ *15 for each group*). **(J)** Fold change in α-SMA stain intensity, normalized to the 0% control group (*n* ≥ *4 for each group*). **(J)** Fold change in lectin stained vessel area, normalized to the 0% control group (*n* ≥ *3 for each group;*). *All data presented as mean* ± *SD*. O*ne-way ANOVA with Tukey’s post-hoc* \**P* < *0*.*05*, \*\**P* < *0*.*01*, \*\*\**P* < *0*.*001*, \*\*\*\**P* < *0*.*0001*

## 4. Discussion

The integration of bioscaffolds with the surrounding host microenvironment is critical for the success of the implant, as poor tissue continuity can hinder cell migration, nutrient transport, and neovascularization, leading to impaired graft functionality (25). It is challenging to optimize this integration since many types of cells are involved in the process, such as monocytes/macrophages, fibroblasts, and endothelial cells (26, 27). Most approaches seek to mitigate the inflammatory response and promote reparative angiogenesis by engineering material properties and localized agent delivery (28, 29); however, the design of such approaches must be carefully tailored to avoid detrimental effects. For instance, elevated dosages of anti-inflammatories can globally suppress macrophage mobility, which impairs host cell integration and vascularization (13, 30). Furthermore, designing a drug delivery platform using a macroporous scaffold is complicated by their low total volume, which decreases drug availability, and their high surface area, which elevates early release kinetics. To enhance the versatility of these platforms for local drug delivery, agents can be integrated into the scaffold using discreet microbeads, as opposed to monolithic dispersions (31-33). In this manner, drug kinetics are more precisely controlled while also permitting for ease in the use of multiple agents.

In this study, PDMS drug-eluting microbeads with an average diameter size of 75 µm were fabricated. Previous studies have employed microfluidics devices to generate PDMS microparticles (18, 34-36); however, the requirement of custom-designed equipment decreases ease in translation across laboratories and scale-up for mass production. Further, for some devices, cleaning PDMS debris post-fabrication is challenging. An alternative method developed for this study used emulsions and a filter-based approach, a 41 µm filter to emulsify the PDMS microbeads and a 100 µm filter for sorting. This method resulted in efficient fabrication using basic laboratory equipment. To further enhance scale-up, larger commercially available meshes could replace the smaller ones used in this study. While our method generated spherical beads with an average size of 75 µm, the high variance in bead size (standard deviation of 51 µm) indicates the coalescence of the microdroplets in the collection solution before PDMS crosslinking. The inclusion of surfactant in the collection solutions, such as sodium dodecyl sulfate (SDS) (34) or Brij® L4 (35), could be used to prevent this issue. Once fabricated, the integration of microbeads into the macroporous scaffold did not impact the scaffold structure or porosity, as validated via microscopy (stereomicroscopy and SEM) images and Nano-CT scanning videos. Thus, these highly porous (85%) scaffolds can be used for both monolithic and microbead drug delivery without impacting the scaffold’s capacity to support robust host cell infiltration and intra-device vascularization, as previously established (2, 13, 37). Investigation of initial drug release from the scaffolds via zero-order kinetic modeling quantified a significant reduction in drug burst release for the microbead format, when compared to the monolithic format; however, the zero-order model was unable to reflect the entire releasing period. To characterize the durability of drug release, the monolithic scaffold was modeled as a monolithic solution matrix system, while the microbead scaffold was modeled as a reservoir device with non-constant activity source (i.e. entirely first-order release) (38). Due to the fact that all MoS groups released 40% of total drugs around day 3 (i.e. 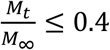), a first-order could be used for analyzing the entire releasing period of MoS. The high goodness-of-fit R^2^ values and the dose-independent first-order release rates for the same scaffold format verified the accuracy of the model. Based on the first-order release model, the durability of drug release was generally extended for µBS groups, when compared to the same drug loading in the MoS format.

The implantation of Dex-eluting scaffolds into the epididymal fat pad of a murine model permitted examination of host responses. Scaffolds were explanted at 14 days post-implantation, as this time point exhibited peak host cell infiltration for drug-free implants (37). For all implants containing Dex, a gradient in infiltrating cells from the scaffold/host interface to the scaffold center was observed, illustrating a spatial gradient in drug release within the implant site. A negative correlation was observed between the degree of host cell infiltration and the dosage of Dex loaded into the scaffold. For both scaffold formats, the 1% and 0.5% Dex loadings were highly suppressive of host infiltration and ECM deposition, indicating an unsuitable dosage for ensuring favorable engraftment of the implant. For lower drug loadings of 0.1% Dex, the significant differences in host cell infiltration between μBS and MoS correlated to the kinetic release curves, indicating a more durable release rate when the same amount of drug was integrated into a microbead versus monolithic format. Suppression of host cell infiltration, ECM deposition, and the formation of vascular structures within the μBS was also observed for the lowest drug loading (0.05%). Based on these observations, Dex exhibited a profoundly suppressive impact on host cell infiltration when delivered using a more durable release format. One approach to mitigate these negative impacts would be to further reduce the Dex loading or use the MoS format; however, decreasing Dex dosage will concurrently decrease its impact on positive macrophage polarization (13, 39). To overcome the negative impacts of local Dex delivery on host cell mobility while retaining its positive features of macrophage polarization, we explored the co-delivery of Dex with the pro-angiogenic agent estrogen (E2).

The translation of our controlled drug delivery system to other agents, i.e. from Dex to E2, was simplified by predictive modeling. Using our structure-related first-order kinetic model and accounting for changes in drug molecular weight, the E2 loading resulting in desired early release rates was successfully estimated. Estrogen release kinetics from μBS were similar to Dex, validating the utility of the system for controlled release of different agents with similar hydrophobicity and solubility. For the lower E2 loading of 0.01%, elevated host cell infiltration and ECM deposition were observed for explants, when compared to the drug-free control. This observation of enhanced cellular activities aligns with published reports that delivered a similar local dosage (5 ng/mL per day), which promoted endothelial cell migration and vasculature formation (16). The positive impact of E2 can be attributed to the local up-regulation of VEGF gene expression via estrogen receptors α and β (15). Unexpectedly, when the E2 dosage in the μBS was increased to 0.1%, inhibition of cellular infiltration and ECM deposition was observed. Others have reported that higher dosages of E2 (i.e. 500 ng/mL) could inhibit vascular smooth muscle cell proliferation and migration (40). Furthermore, E2 reduces the expression of membrane E-selectin (41). The global effect of E2 on host responses and inflammation is complex, with reports of dose-dependent impacts on various cell types (42). With the unique dose-dependency observed in this study, further investigation into the role of E2 and scaffold engraftment is needed to identify therapeutic local doses.

The incorporation of dual drug microbeads, Dex and E2, resulted in a local, dual-drug delivery system. The multi-drug release profiles could also be described by the structure-related first-order model, supporting the versatility of this microbead scaffold approach to deliver multiple drugs with predictable release profiles that are minimally impacted by each other. Cytokine profiling of LPS-stimulated M1 macrophages co-cultured with E2 and/or Dex eluding scaffolds provided insight into the role of each agent in directing macrophage phenotype. As expected, Dex-only scaffolds reduced the secretion of pro-inflammatory cytokines IL-6 and TNFα, as well as pro-angiogenic VEGF (13, 43-45). While LPS stimulation of macrophages can inherently stimulate IL-10 production as a part of a negative feedback loop, Dex damped this response (46, 47). The co-culture of stimulated macrophages with E2-only scaffolds did not appear to impart any significant effects on the measured cytokines. Previous studies observed a down-regulation in IL-6 and TNFα expression when murine macrophages were pre-treated with E2 (0.1 ng/mL) (48); however, our scaffolds were added post-LPS treatment. The combinatory impact of E2 and Dex on macrophage phenotype, per this specific cytokine screening, indicates that Dex has the most dominant effect. The inclusion of E2 with the Dex did not significantly alter IL-6 or TNFα release, although it did boost IL-10 and VEGF secretion. These results support the capacity of E2 to elevate the pro-angiogenic phenotype of macrophages when they are co-exposed to Dex.

To investigate this theory *in vivo*, dual-delivery scaffold implants were tested in a murine model. While systemic impacts on circulating blood levels were not observed, remarkable impacts on host cell remodeling at the local implant site were measured. The E2-only group displayed elevated matrix deposition, α-SMA deposition, and functional vasculature within the scaffold implant, when compared to the control group. As previously mentioned, E2’s positive impacts on endothelial cells and fibroblasts could contribute to these effects (16, 42). The Dex-only group displayed expected suppression in infiltrated cells, α-SMA deposition, and functional vasculature within the scaffolds, when compared to drug-free implants. Results imply that a low Dex loading of 0.05% from the μBS format still had significant impacts on mitigating cellular migration, which resulted in an undesired delay in vasculature formation and/or competency. The combination of local E2 and Dex release from the μBS resulted in unique host responses, with a significant increase in infiltrated cell number and matrix deposition, when compared to the Dex-only group. These observations indicate E2 counter-regulates Dex effects on migration. When compared to the E2-only μBS explants, Dex+E2-μBS scaffolds exhibited depressed cellular infiltration, ECM deposition, and vascularization, indicating Dex also counters E2 effects. When compared to drug-free implants, the combination scaffold exhibited an overall suppressive impact on the cellular infiltration, while contrarily boosting α-SMA and maintaining competent vascularization. Based on these results, the combination of 0.01% E2 and 0.05% Dex has the potential to create a favorable host environment, where the counter-regulation of E2 and Dex results in a robustly engrafted implant. Furthermore, it highlights the potential of this multi-drug delivery microbead scaffold format to create customized host responses to the local microenvironment.

Future studies will focus on further optimizing the concentration of drugs to understand more about the drug release model and the combined effects of Dex and E2. Furthermore, the local impact of these agents on specific host responses will also be parsed out via delineation of cell types and characterization of phenotypes. The approach will also be translated to different agents and combinations to explore their local effects. Finally, the Dex/E2 dual drug delivery system will be evaluated in cellular transplant models to investigate their potential benefits in graft efficacy.

## 5. Conclusion

Herein, a distinct, versatile, dual drug delivery system based on a microbead integration into a macroporous scaffold was developed. To fabricate the PDMS microbeads, a simple and efficient method was developed, which could be easily translated for mass production. This system showed the potential to modulate burst release and extended the durability of drug release, while simultaneously permitting ease in dual drug release. *In vivo* studies correlated with drug kinetic studies, demonstrating a means to design a predictable system that translates *in vivo*. The combination of two complementary drugs, Dex and E2, imparting unique effects on host cell responses, providing a balance in host cell infiltration and vascularization. Future studies will seek to leverage this platform for the controlled release of numerous other agents, as well as exploring the potential of using the Dex+E2 scaffold system to create a favorable microenvironment for cellular transplantation.

## Acknowledgments

We thank Irayme Labrada-Miravet for her excellent technical assistance during animal transplantation. We thank Sydney Wiggins for her assistance in isolation and differentiation of bone-marrow derived macrophages. We thank all members of Stabler Laboratory for their collective assistance in animal care and monitoring. We thank Electron Microscopy Core of Interdisciplinary Center for Biotechnology Research, Research Service Centers, and Sharma Laboratory at the University of Florida for equipment use. We thank Molecular Pathology Core at the University of Florida for assistance in processing histological samples.

## SUPPLEMENTARY MATERIALS

**Figure S1.**
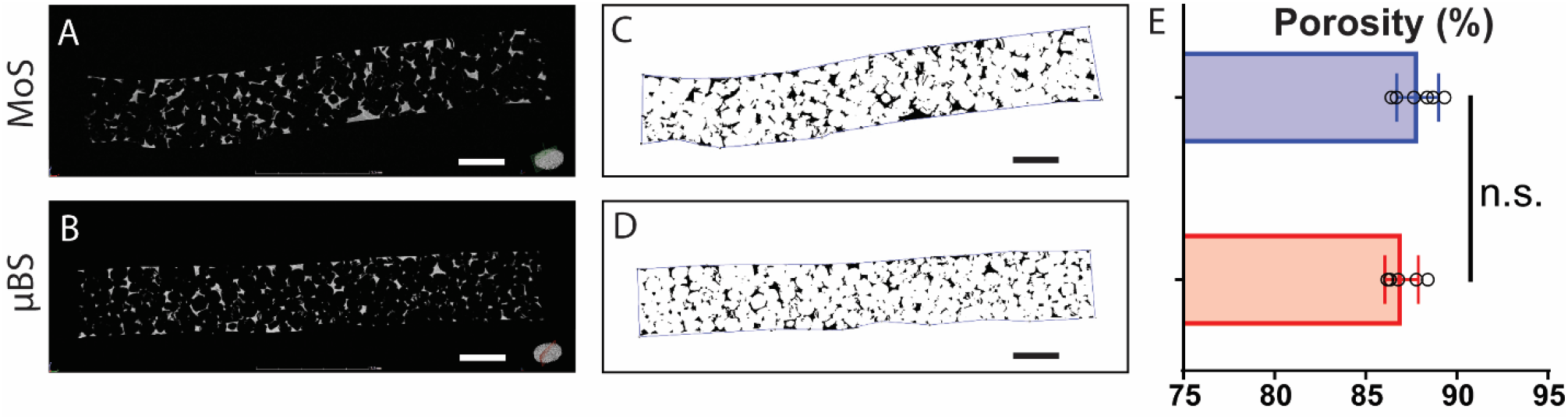
NanoCT visualization and quantification of porosity for monolithic and microbead scaffold formats. **(A-B)** Representative NanoCT scanning images of the scaffold cross-section, illustrating porosity and pore interconnectivity of the scaffolds. **(C-D)** Binary processed images used for porosity quantification. **(E)** Comparison of % porosity between the two scaffold formats (*data sourced from 6 different cross-sectional processed NanoCT images; data presented as mean* ± *SD; P* > *0*.*05, not significant, unpaired t test*). *Scale bar = 1 mm*.

**Figure S2.**
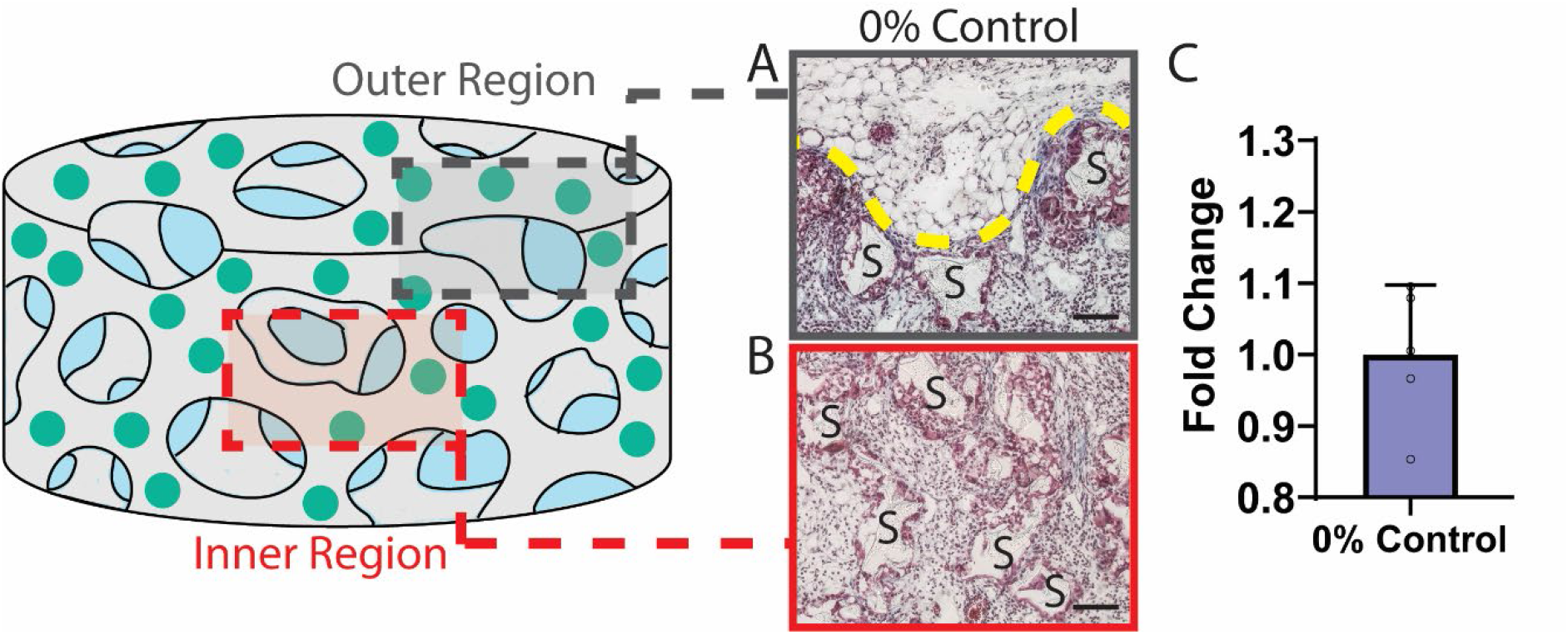
Trichrome staining of drug-free scaffolds explanted from epididymal fat pad 14 days post-implantation. **(A)** Trichrome stained cross-section of control scaffold collected from the host/scaffold interface (designated by schematic as the “Outer Region”). *Yellow dashed line = scaffold/host interface* **(B)** Trichrome stained cross-section of control scaffold collected from the center of the scaffold implant (designated by schematic as the “Inner Region”). **(C)** Fold change in nuclei counts, as quantified from processed images of the Inner Region and normalized to the 0% control group (*n = 5; data presented as mean* ± *SD*). *“S” = PDMS scaffold. Scale bar = 100 μm*.

**Figure S3.**
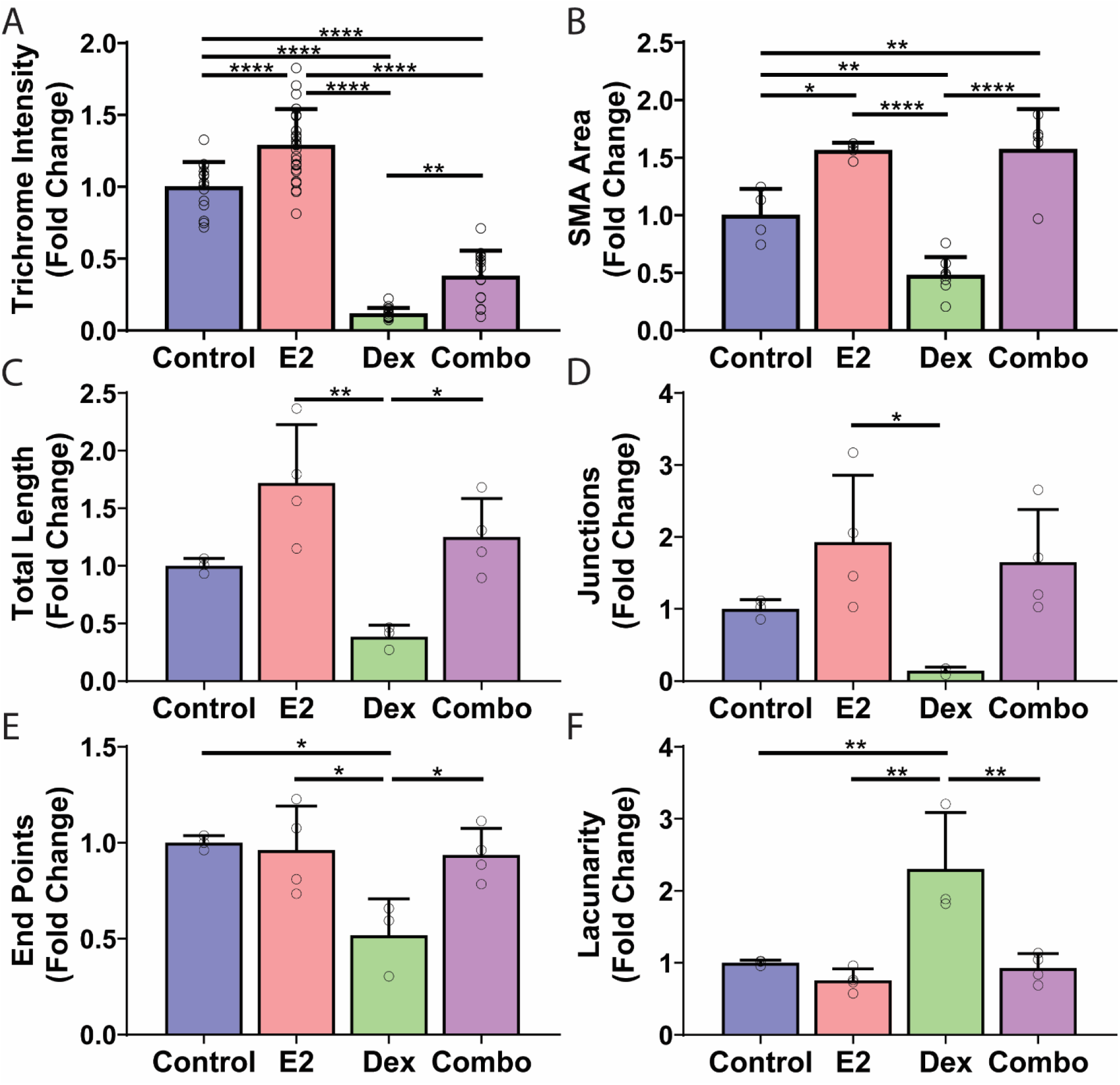
Supplemental analyses of epididymal fat pad explants of drug-loaded μBS retrieved 10 days post-implantation. **(A)** Trichrome staining intensity normalized to the control group (*n* ≥ *15 for each group*). **(B)** α-SMA staining area normalized to the control group (*n* ≥ *4 for each group*). **(C-F)** Total vessel length, total number of junctions, total number of end points, and lacunarity of perfused blood vessels (*n* ≥ *3 for each group*). *All data presented as mean* ± *SD. Statistical analysis: One-way ANOVA with Tukey’s multiple comparisons test* \**P* < *0*.*05*, \*\**P* < *0*.*01*, \*\*\*\**P* < *0*.*0001*

**Figure S4.**
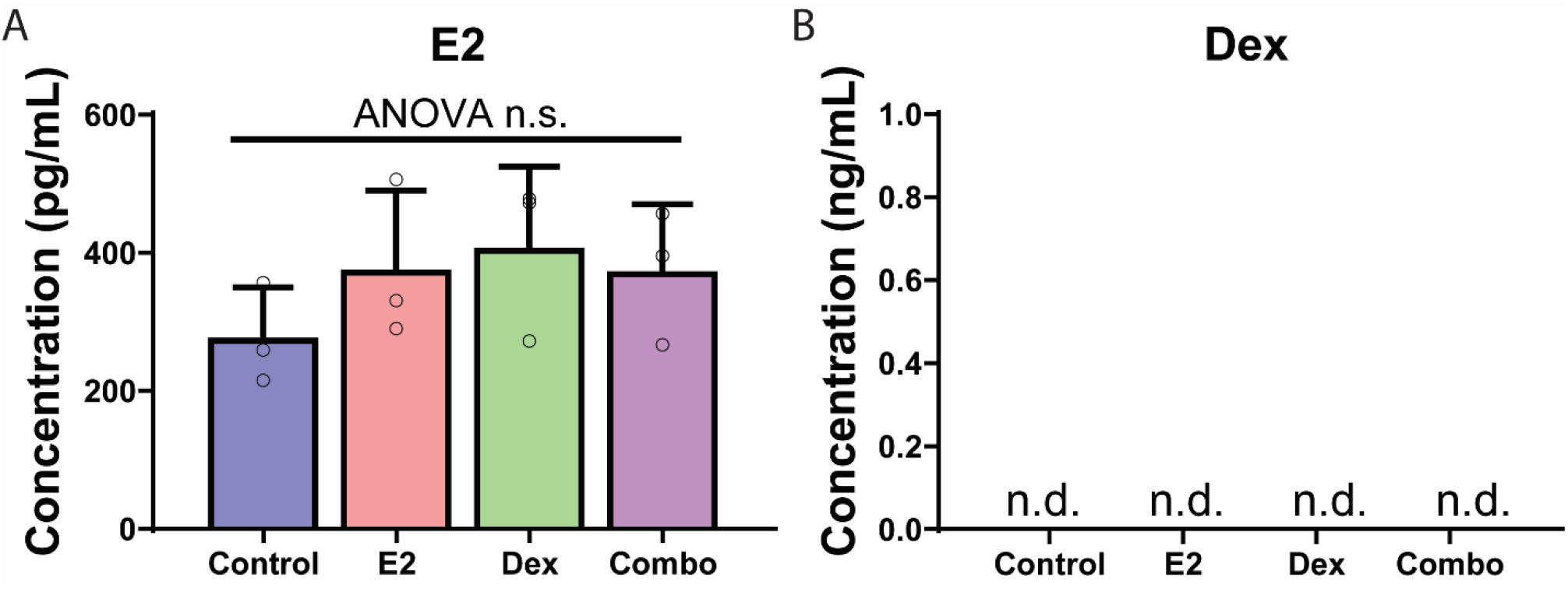
Circulating plasma levels of estrogen and dexamethasone 10 days post-implantation of designed µBS. **(A)** E2 concentration in circulating plasma (*n = 3 for each group; data presented as mean* ± *SD; P* > *0*.*05, not significantly different, one-way ANOVA with Tukey’s multiple comparisons test; n*.*s. = not significant*). **(B)** Dex concentration in circulating plasma (*n = 3 for each group; not detectable; ELISA kit sensitivity = 0*.*23 ng/mL*).

**Table S1.** Summary of First Order Release Analysis of Dex Eluting Scaffolds

| Scaffold Type | Zero Order |  | First Order |  |  |  |
| --- | --- | --- | --- | --- | --- | --- |
| | Rate (ng/day) | R<sup>2</sup> | Rate (1/day) | Half-time | Half-time (μBS/MoS) | R<sup>2</sup> |
| 1% MoS | 2146 | 0.95 | 0.143 | 4.83 | — | 0.91 |
| 1% μBS | 327.6 | 0.97 | 0.084 | 8.25 | 1.7 | 0.99 |
| 0.5% MoS | 717.0 | 0.89 | 0.210 | 3.30 | — | 0.85 |
| 0.5% μBS | 128.2 | 0.98 | 0.022 | 31.8 | 9.6 | 0.99 |
| 0.25% MoS | 53.82 | 0.95 | 0.240 | 2.89 | — | 0.98 |
| 0.25% μBS | 17.95 | 0.93 | 0.107 | 6.49 | 2.2 | 0.94 |
| 0.1% MoS | 7.405 | 0.76 | 0.175 | 3.96 | — | 0.71 |
| 0.1% μBS | 11.07 | 0.89 | 0.112 | 6.16 | 1.6 | 0.92 |
| 0.05% μBS | 5.174 | 0.97 | 0.103 | 6.73 | — | 0.97 |

### Supplementary Text 1. E2 Concentration Estimated from Dex Release Profile

A positive pro-angiogenic effect of local E2 was reported at the first-day release of ∼5 ng E2/day (1). To select an E2 dosage within the μBS that would generate this desired release, modeling theory and extracted parameters collected from Dex release curves were extrapolated to predict release profiles for E2-μBS. The first order diffusion-controlled release model for drug release from the PDMS macroporous scaffold (based on (2)) was defined as:

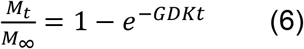

where *M*_*t*_ (ng) is the cumulative amount of drug released at time t (day), *M*_0_ (ng) is the initial amount of drug in the solution, *M*_∞_ (ng) is the total amount of drug that could be released from the scaffolds, G is a ratio related to the scaffold geometry, D is the diffusion coefficient of the drug within PDMS, and K is the partition coefficient of the drug between the PDMS microbead and its surrounding environment.

The first-day fractional release can be calculated by reducing equation (6) to

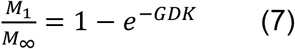

Assuming that the scaffold geometry and PDMS material properties were not impacted by the drug type and comparable solubility of Dex and E2 (further mitigated by the low drug loading), the parameters G and K should not significantly change when replacing the drug Dex with E2; therefore, the first-day fractional release of scaffolds was only directly impacted by D.

Malcolm et al. reported a linear relationship between the logarithm of the D (cm^2^/s) and the molecular weight of the drug (MW; g/mol) (3) in PDMS, specifically:

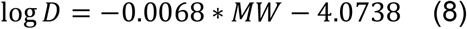

Examination of Dex release curves found the 0.05% wt/wt dosage to exhibit a release rate in the desired E2 range, thus modeling parameters sourced from this μBS dosage were used (**Table S2**). Designating *M*_1_ for the E2-μBS to be 5 ng, *M*_∞_ for the E2-μBS could be derived from equation (7) and (8), and used to estimate the desired drug concentration for the μBS (**Table S2**). The predicting E2 concentration needed to reach the targeted 5 ng first-day release rate was 0.007% wt/wt μBS, which was rounded up to 0.01% wt/wt E2 for ease in experimental accuracy.

**Table S2.** Prediction of E2 Drug Loading into μBS using Models Generated from 0.05% Dex-μBS Experimental Data

| Drug Type | MW (g/mol) | log D ($\text{cm}^2/\text{s}$) | D ($\text{cm}^2/\text{s}$) | D(Dex)/D(E2) | First Day Fractional Release (%) | First Day Release (ng) | Concentration (% wt/wt) |
| --- | --- | --- | --- | --- | --- | --- | --- |
| Dex | 392.461 | -6.74 | 1.81E-07 | — | 0.12 | 7.7 | 0.05 |
| E2 | 272.38 | -5.93 | 1.19E-06 | 6.6 | 0.57 | 5.0 | 0.01 |

**Figure S5.**
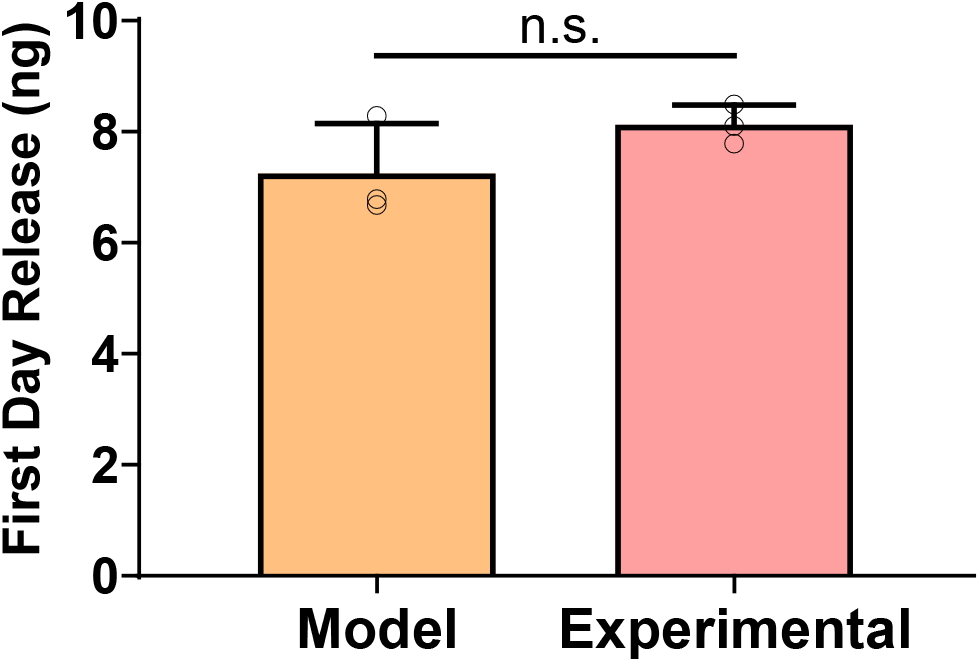
First Day Release from 0.01% wt/wt E2 Scaffolds. (P > 0.05, not significant, unpaired t test).

**Table S3.**
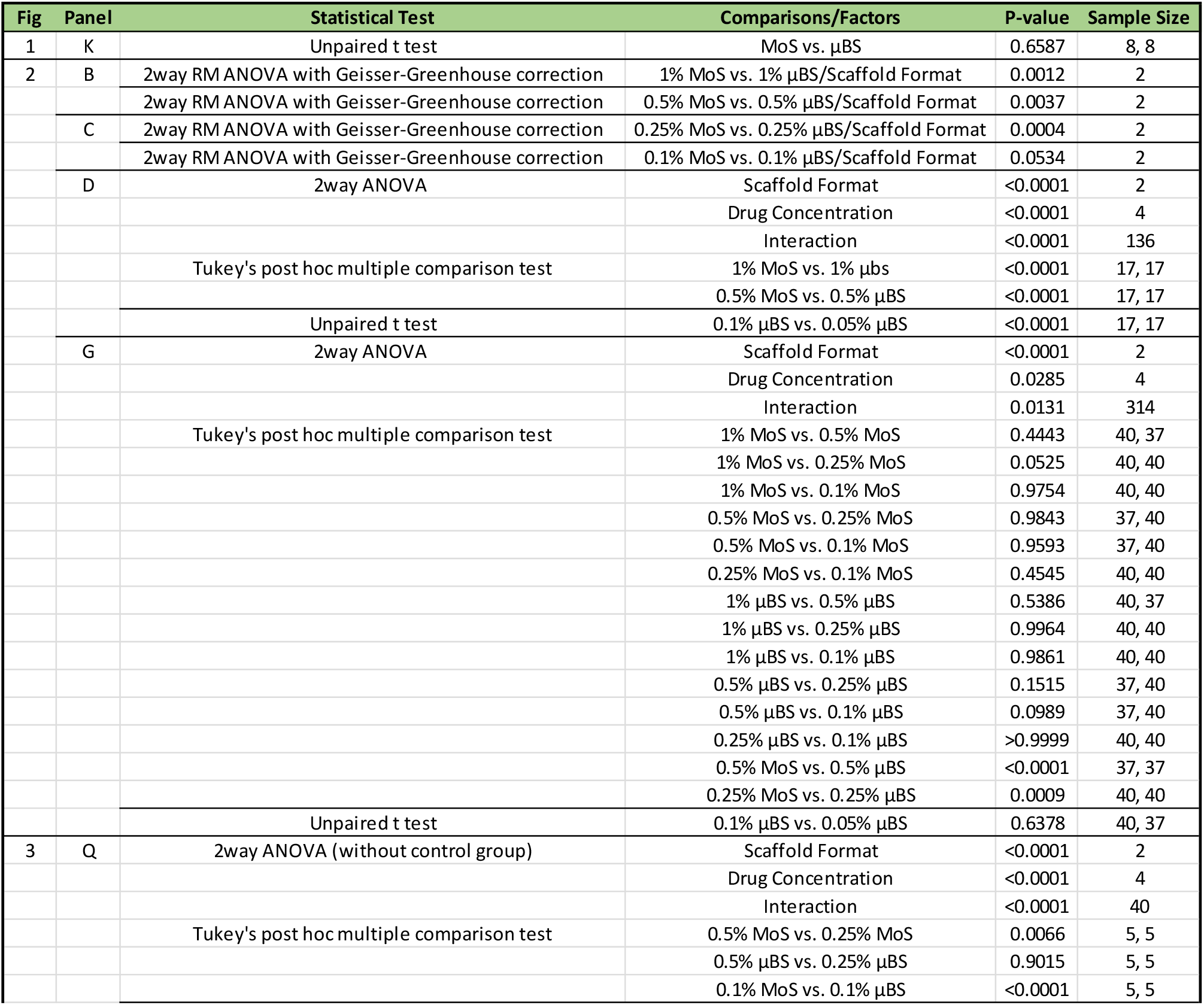
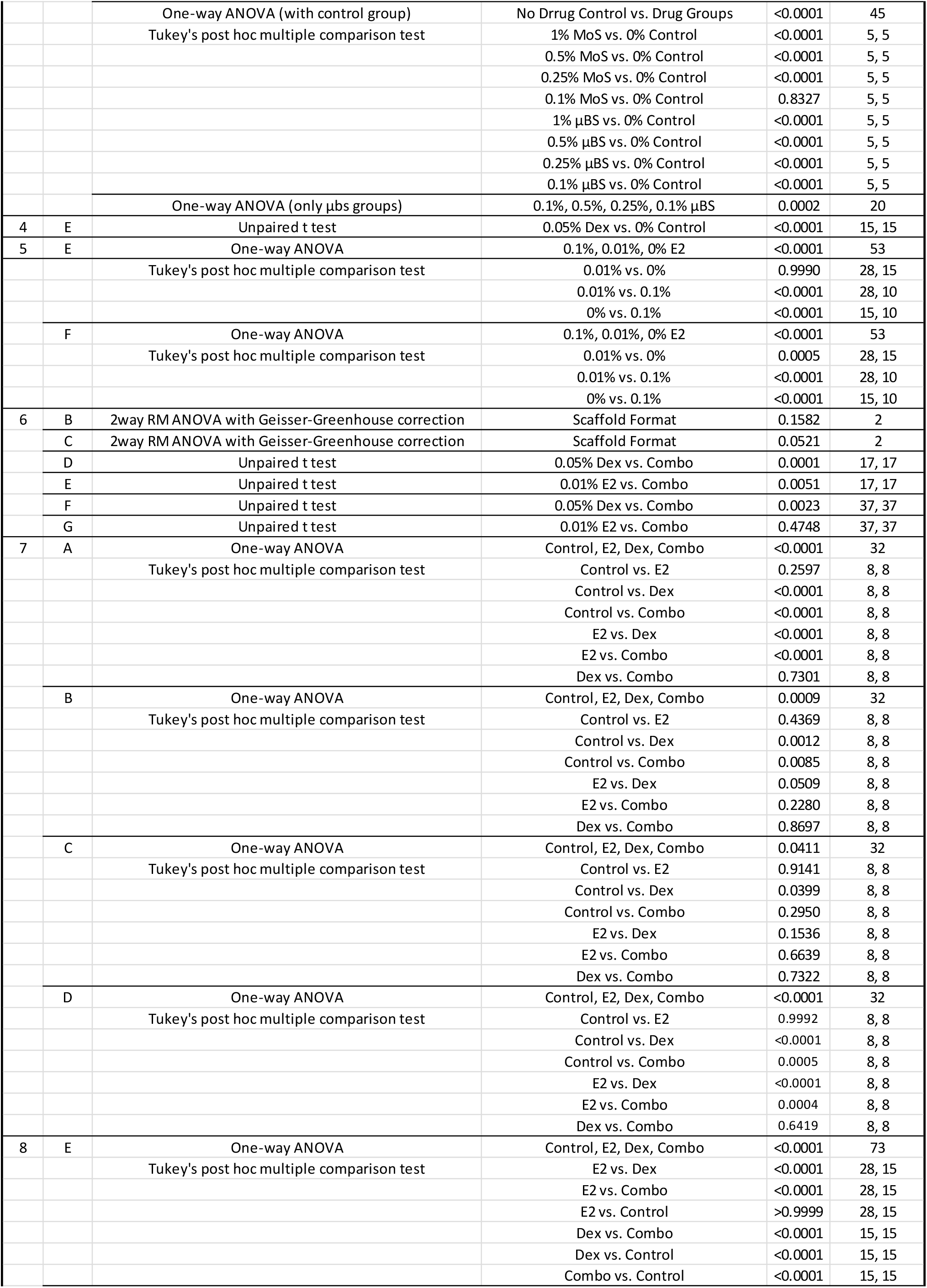
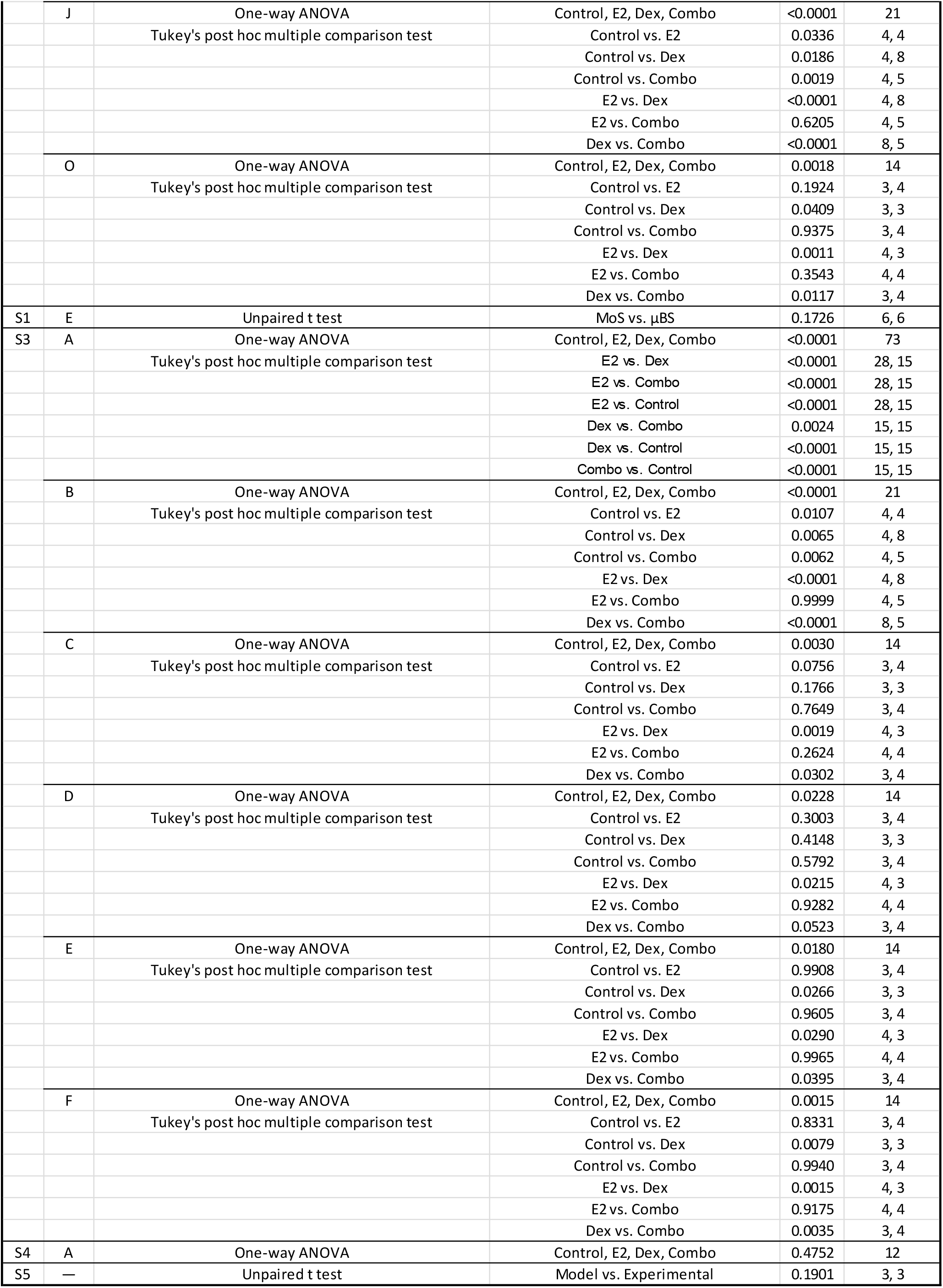
Full Summary of Statistical analysis.

